# Cell wall integrity and elicitor peptide signaling modulate jasmonic acid-mediated camalexin production in Arabidopsis

**DOI:** 10.1101/2025.09.09.675042

**Authors:** Richard Noi Morton, Lian Fleischberger, Joy Debnath, David Biermann, Susanne Mühlbauer, Ahalya Rajendran, Hans-Henning Kunz, Julien Gronnier, Nora Gigli Bisceglia, Timo Engelsdorf

## Abstract

Plant cell walls constitute dynamic barriers that are essential for defense against pathogens. The receptor kinase THESEUS1 (THE1) monitors cell wall integrity (CWI) and contributes to pathogen resistance in Arabidopsis, but the underlying mechanisms remain unclear. Here we show that THE1-dependent CWI signaling induces accumulation of the antimicrobial metabolite camalexin upon cell wall damage (CWD) caused by cellulose biosynthesis inhibition or fungal infection. CWD alters THE1 plasma membrane nanodomain organization and involves calcium signaling components that modulate camalexin production. Induction of camalexin requires jasmonic acid (JA)-dependent expression of the transcription factors MYB47 and MYB95. In line with its antagonistic function on CWI signaling, the plant elicitor peptide Pep3 suppresses camalexin biosynthesis downstream of THE1 by inhibiting JA-dependent pathways. Our findings reveal a regulatory network where CWI and Pep3 signaling modulate antimicrobial defense via JA-mediated camalexin production. This network requires independent CWD-induced pathways, providing insights into how plants balance defense activation and suppression in response to cell wall stress.

## Introduction

Cell walls provide mechanical support to plant cells and serve as sturdy, yet highly dynamic barriers against plant pathogen infection. The ability of pathogens to modify host cell wall properties and to locally breach the barrier determines their successful entry into host tissues and their virulence on specific host plants (Bhandari et al., 2025; Munzert and Engelsdorf, 2025). Consequently, specific alterations in cell wall composition and structure influence the outcome of plant-pathogen interactions (Molina et al., 2021; Reckleben et al., 2025). It is well established that immune signaling is increased in many cell wall mutants and accumulating evidence suggests that signaling pathways induced by cell wall damage (CWD) contribute to pathogen defense (Bacete et al., 2018; Vaahtera et al., 2019). The cell wall status is constantly surveilled by a number of overlapping processes together referred to as cell wall signaling (Wolf, 2022). Cell wall stress correlates with the expression of small signaling peptides, which can influence cell wall properties (Debnath et al., 2026). Receptor kinases of the *Catharanthus roseus* receptor-like kinase 1-like (*Cr*RLK1L) family have a prominent role in cell wall signaling during development, growth, damage and immunity (Franck et al., 2018; Ortiz-Morea et al., 2022; Wolf, 2022). *Cr*RLK1Ls are receptors for RAPID ALKALINIZATION FACTOR (RALF) family peptides, which are central regulators of *Cr*RLK1L signaling (Schoenaers and Vissenberg, 2025). The *Cr*RLK1L THESEUS1 (THE1) is required for the induction of responses to CWD, e.g. in cellulose-deficient mutants, after cellulose biosynthesis inhibition with isoxaben (ISX) or after enzymatic cell wall degradation (Hématy et al., 2007; Engelsdorf et al., 2018; Bacete et al., 2022). Binding of RALF34 to THE1 contributes to regulation of lateral root initiation but is dispensable for THE1-dependent responses to ISX (Gonneau et al., 2018). Signaling responses induced by CWD are referred to as cell wall integrity (CWI) signaling (Vaahtera et al., 2019; Wolf, 2022). Responses to ISX include the reactive oxygen species (ROS)- and calcium-dependent induction of phytohormone production and ectopic cell wall lignification (Hamann et al., 2009; Denness et al., 2011). These responses depend on THE1 and the receptor kinases MALE DISCOVERER 1-INTERACTING RECEPTOR LIKE KINASE 2 (MIK2) and STRUBBELIG (Van der Does et al., 2017; Chaudhary et al., 2020). THE1 and MIK2 have partially overlapping functions in CWI signaling and both contribute to defense against the soilborne fungus *Fusarium oxysporum* (Van der Does et al., 2017). MIK2 is the receptor for SERINE-RICH ENDOGENOUS PEPTIDEs (SCOOPs) that regulate immunity and senescence (Hou et al., 2021; Rhodes et al., 2021; Zhang et al., 2024). ISX-induced CWD promotes SCOOP expression in a partially THE1-dependent manner, which increases responses to a subsequent treatment with pathogen-associated molecular patterns (PAMPs) (Zhai et al., 2024). This highlights how signals originating from a damaged cell wall can contribute to pattern-triggered immunity (PTI). Intriguingly, loss of PTI elements such as BRASSINOSTEROID INSENSITIVE 1 (BRI1)–ASSOCIATED KINASE 1 (BAK1), BAK1-LIKE 1 (BKK1), BOTRYTIS-INDUCED KINASE 1 (BIK1), PEP1 RECEPTOR 1 (PEPR1) and PEPR2 leads to increased ISX responses, indicating that PTI can negatively regulate CWI signaling (Engelsdorf et al., 2018).

PTI’s feedback inhibition of CWI signaling depends on endogenous plant elicitor peptides (Peps) that are perceived by PEPR1/2 and BAK1 (Yamaguchi et al., 2006; Yamaguchi et al., 2010; Tang et al., 2015). Pep1 and Pep3 precursor genes, *PROPEP1* and *PROPEP3*, are induced in a THE1-independent manner after 1 hour of ISX treatment and co-treatment of seedlings with ISX and synthetic Pep1 or Pep3 suppresses ISX-induced accumulation of the phytohormones jasmonic acid (JA) and salicylic acid (SA) (Engelsdorf et al., 2018). *PROPEPs* are also induced by wounding, salinity, pathogen infection or PAMP treatments, but the direct stimulus leading to their expression is unknown (Bartels and Boller, 2015; Nakaminami et al., 2018). Salinity-induced *PROPEP3* expression is alleviated by treatments that affect cell wall crosslinking, indicating that stress-induced changes in cell wall properties may play a role (Gigli-Bisceglia et al., 2022). Overexpression of *PROPEP1* and *PROPEP3* is sufficient to rescue plants from cell death after long-term treatment with ISX or growth under high salinity, demonstrating their impact on stress resistance (Nakaminami et al., 2018; Zhang et al., 2025). Cellular damage activates calcium-, SUMOylation- and metacaspase-dependent PROPEP processing and released Peps can spread danger signals to neighboring cells (Hander et al., 2019; Shen et al., 2019; Zhang et al., 2025).

In summary, the state of the art suggests that THE1-dependent CWI monitoring can contribute to basal defense against pathogen infection (Guerreiro and Marhavý, 2023). Basal defense against pathogens involves the production of phytohormones and of specialized antimicrobial metabolites, termed phytoalexins (Bari and Jones, 2009; Ahuja et al., 2012). In Arabidopsis, the major phytoalexin camalexin is synthesized by a metabolon comprising several Cytochrome P450 enzymes, whose expression is regulated by WRKY transcription factors (Birkenbihl et al., 2012; Birkenbihl et al., 2017; Mucha et al., 2019). During fungal infections, camalexin is produced and secreted locally at infection sites to restrict fungal proliferation (Glawischnig, 2007; He et al., 2019).

Here, we investigated the mechanism by which camalexin biosynthesis is activated and regulated upon CWD through THE1-dependent CWI signaling and how this defense response is negatively regulated by Pep3 signaling. Our findings elucidate the interplay between these pathways in modulating antimicrobial defense in Arabidopsis.

## Results

### THE1 contributes to camalexin accumulation and pathogen resistance

To investigate how THE1-mediated CWI signaling contributes to pathogen resistance, we infected the *the1-1* loss of function mutant with the hemibiotrophic fungus *Colletotrichum higginsianum*, which depends on host cell wall penetration, causing local CWD (O’Connell et al., 2004; Engelsdorf et al., 2017). The fungal entry rate was significantly increased in *the1-1*, indicating that CWI signaling contributes to defense at this early infection step (Figure 1A). To identify antimicrobial defense responses linked to CWD, we examined publicly available transcriptomics data and found that genes encoding for Cytochrome P450 enzymes involved in camalexin biosynthesis are strongly upregulated after ISX-induced CWD (Hamann et al., 2009; Figure S1). Previous research demonstrated that camalexin-deficiency caused by loss of *CYP71B15/PAD3* or *WRKY33*, a major positive regulator of camalexin biosynthesis, critically weakens resistance against *C. higginsianum* (Narusaka et al., 2004; Mao et al., 2011; Schmidt et al., 2020). Indeed, camalexin accumulation was reduced in *the1-1* after *C. higginsianum* infection, indicating that THE1 contributes to induction of camalexin biosynthesis upon CWD caused by fungal infection (Figure 1B). To further investigate the importance of impaired camalexin accumulation for *the1-1* susceptibility, we crossed *the1-1* with the camalexin-deficient *pad3* mutant and quantified fungal genomic DNA during the necrotrophic growth phase, when colonization of surrounding tissue is increased in *pad3* (Narusaka et al., 2004). Fungal DNA was significantly increased in *the1-1*, *pad3* and *the1-1 pad3* compared to Col-0 wild type, but not significantly different between single and double mutants, indicating overlapping effects of THE1 and camalexin synthesis on pathogen resistance (Figure 1C).

**Figure 1:**
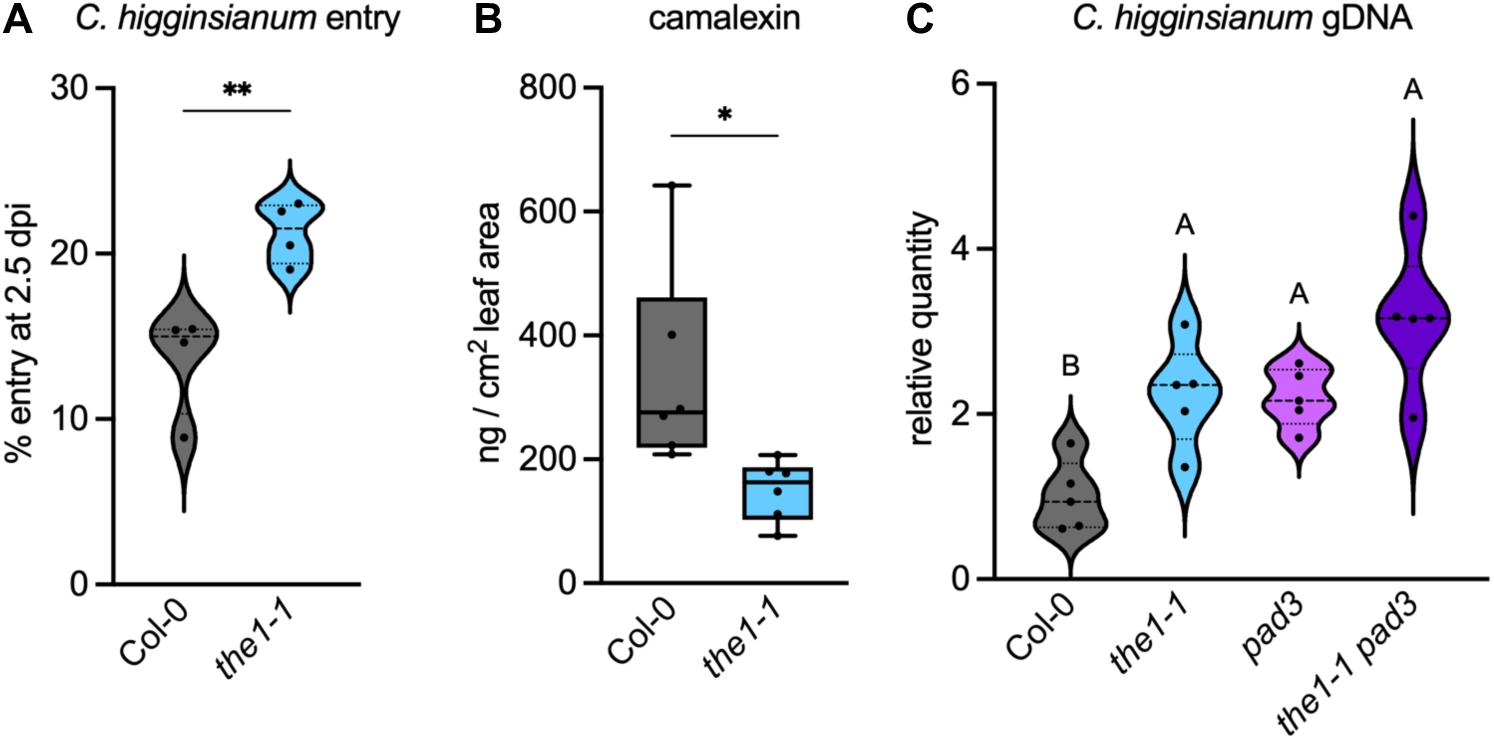
THE1 and camalexin biosynthesis have overlapping roles in resistance to *Colletotrichum higginsianum* infection. (A) Entry rate of *C. higginsianum* on Col-0 and *the1-1* leaves at 2.5 days post infection (dpi) (n=4). (B) Camalexin amount in infected Col-0 and *the1-1* leaves at 2.5 dpi (n=6). Asterisks in (A,B) indicate statistically significant differences to Col-0 according to a Student’s t-test (*p < 0.05, **p < 0.01). (C) The relative quantity of *C. higginsianum* DNA was determined at 4 dpi in Col-0, *the1-1*, *pad3* and *the1-1 pad3* (n=5). Different letters indicate statistically significant differences according to one-way ANOVA and Holm-Sidak’s multiple comparisons test (α = 0.05).

### Cell wall damage is sufficient to induce THE1-dependent camalexin accumulation

The finding that THE1 played a role in camalexin induction during pathogen entry through the cell wall prompted us to investigate if CWI impairment is sufficient to induce camalexin biosynthesis. ISX treatments are ideally suited to further investigate the regulation of CWD-induced camalexin biosynthesis, as treatments can be performed in seedling liquid culture in a controlled and well-established manner separating CWD from other molecular events that occur during plant-pathogen interactions (Paredez et al., 2006; Gutierrez et al., 2009). Consistent with transcriptional activation of camalexin biosynthesis, ISX treatments of 10 days old seedlings induced camalexin accumulation after 8 hours (Figure 2A). We have previously shown that treatment of Arabidopsis seedlings with Driselase, a mixture of cell wall-degrading enzymes, induces similar CWI responses as ISX treatment (Engelsdorf et al., 2018). In line with that, camalexin accumulation was increased after Driselase treatment compared to heat-inactivated Driselase, indicating that camalexin accumulation is induced by different types of CWD (Figure S2). Stress-induced conversion of the key indolic intermediate indole-3-acetaldoxime (IAOx) to indole-3-acetonitrile (IAN) for camalexin biosynthesis depends on the two cytochrome P450 enzymes CYP71A12 and CYP71A13, with stronger contribution of the latter (Müller et al., 2015). Transcriptional data indicated that *CYP71A12* was already induced after 4 hours ISX treatment, whereas *CYP71A13* was upregulated 8 hours after ISX treatment, following a similar expression pattern as *CYP71B15/PAD3*, the final enzyme in camalexin biosynthesis (Figure S1). Loss of CYP71A13 completely abolished ISX-induced camalexin accumulation, while loss of CYP71A12 increased camalexin accumulation (Figure 2B). This data shows that CYP71A13 is essential for ISX-induced camalexin biosynthesis, whereas CYP71A12 rather diverges IAOx into other pathways such as indole-3-carboxylic acid (ICA) synthesis (Pastorczyk et al., 2020). To investigate if CWD-induced camalexin biosynthesis depends on THE1, we quantified camalexin in ISX-treated *the1-1* seedlings and found that camalexin accumulation was completely abolished in the mutant (Figure 2C). Consistently, treatment of the hypermorphic *the1-4* allele (Merz et al., 2017) caused increased camalexin accumulation (Figure 2C). Camalexin accumulation was abolished in an ISX-treated *wrky33* mutant and close to the detection limit in *the1-4 wrky33* double mutants, showing that the major transcriptional regulator of camalexin biosynthesis, WRKY33, is necessary for ISX-induced, THE1-dependent camalexin synthesis (Figure 2D).

**Figure 2:**
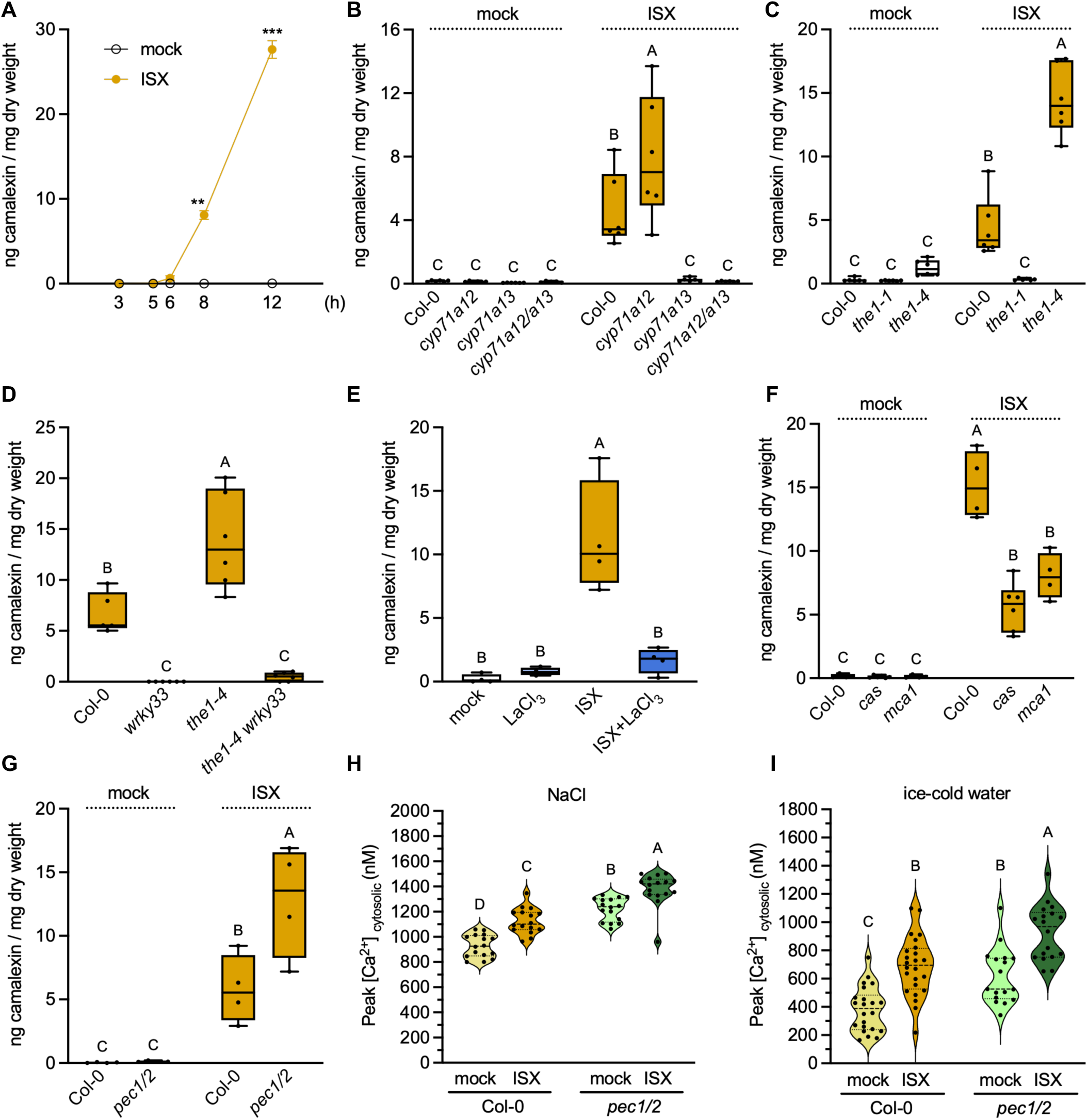
Isoxaben induces camalexin accumulation via THESEUS1, WRKY33 and calcium signaling. (A) Camalexin amount in Col-0 seedlings was determined at 3, 5, 6, 8 and 12 h after mock and isoxaben (ISX) treatment (n=3). Error bars indicate the SEM. Asterisks indicate statistically significant differences between ISX and mock according to a Student’s t-test (*p < 0.05, **p < 0.01, ***p < 0.001). (B-G) Camalexin amount in (B) Col-0, *cyp71a12*, *cyp71a13* and *cyp71a12*/*a13* at 8 h after mock and ISX (n=5-6); (C) Col-0, *the1-1* and *the1-4* at 8 h after mock and ISX (n=6); (D) Col-0, *wrky33*, *the1-4* and *the1-4 wrky33* at 8 h after ISX (n=5-6); (E) Col-0 at 8 h after mock, LaCl_3_, ISX and ISX+LaCl_3_ (n=4); (F) Col-0, *cas* and *mca1* at 8 h after mock and ISX (n=4-6) and (G) Col-0 and *pec1/2* at 8 h after mock and ISX treatment (n=4). (H,I) Peak cytosolic Ca^2+^ values of transients in Col-0 and *pec1/2* seedlings after elicitation with (H) 200 mM NaCl (n=16) and (I) ice-cold water (n=17-24). Prior to elicitation, seedlings were treated for 4.5 h with mock or ISX. Different letters indicate statistically significant differences according to (D,E) one-way ANOVA or (B,C,F,G,H,I) two-way ANOVA and Holm-Sidak’s multiple comparisons test (α = 0.05).

### Calcium signaling determines the magnitude of ISX-induced camalexin response

Previous studies demonstrated that Ca^2+^ signaling is required for ISX-induced JA accumulation (Denness et al., 2011) and that Ca^2+^ transients are involved in CWI maintenance under high salinity (Feng et al., 2018). Consistently, blocking of calcium channels with LaCl_3_ during ISX treatments indicated that Ca^2+^ signaling is required for ISX-induced camalexin accumulation (Figure 2E). To resolve which channels and Ca^2+^ sensing mechanisms contribute to this response, we investigated mutants for the plasma membrane localized channel MATING PHEROMONE INDUCED DEATH 1 (MID1)–COMPLEMENTING ACTIVITY 1 (MCA1), which has been previously implicated in mechanosensing and CWI signaling (Nakagawa et al., 2007; Engelsdorf et al., 2018), and the chloroplast localized CALCIUM SENSING RECEPTOR (CAS), which is required for the regulation of cytosolic Ca^2+^ transients upon external Ca^2+^ signals (Nomura et al., 2008; Weinl et al., 2008). Camalexin accumulation in response to ISX treatment was reduced by 60 and 47 % in *cas* and *mca1* mutants, respectively, supporting that both genes contribute to CWD responses (Figure 2F). Stress-induced Ca^2+^ transients occur in the cytosol and in the chloroplast stroma (Nomura et al., 2012). To test whether altered distribution between stromal and cytosolic Ca^2+^ influences ISX responses, we investigated camalexin production in double mutants of the PLASTID ENVELOPE ION CHANNELs (PEC1, PEC2) that exhibit reduced stromal and increased cytosolic Ca^2+^ transients after stress exposure (Völkner et al., 2021). ISX treatment induced higher camalexin accumulation in *pec1 pec2* than in Col-0, indicating that increased cytosolic Ca^2+^ transients promote the CWD response (Figure 2G). The importance of Ca^2+^ at a later stage (hours) of CWD might indicate that Ca^2+^ signaling is influenced by CWI status. To test this idea, we used high salinity and ice-cold water, two independent stimuli inducing cell wall stress (Debnath et al., 2026), to elicit cytosolic Ca^2+^ transients in Col-0 and *pec1 pec2* seedlings that had been pretreated with mock or ISX. Both treatments caused increased cytosolic Ca^2+^ peaks in ISX-compared to mock-treated seedlings, indicating that CWD increases the sensitivity for Ca^2+^ elicitation (Figure 2H,I; Figure S3).

### Pep3-triggered signaling inhibits CWI signaling downstream of THE1

We have previously reported that ISX-induced, THE1-dependent phytohormone responses are negatively regulated by Pep1 and Pep3, however the underlying mechanism remained unknown (Engelsdorf et al., 2018). To investigate how Pep perception affects ISX-induced camalexin accumulation, we treated Col-0, *pepr1 pepr2* and *bak1-5 bkk1* seedlings with ISX, Pep3 and a combination of both. We chose Pep3 for this study, (i) because it attenuates THE1-dependent phytohormone accumulation, but, unlike Pep1, does not induce ectopic root lignification, and (ii) because we previously demonstrated that PROPEP3 is secreted upon cell wall damage (Engelsdorf et al., 2018). Pep3 co-treatment led to suppression of ISX-induced camalexin accumulation in a PEPR- and BAK1/BKK1-dependent manner (Figure 3A), indicating that Pep3 inhibits THE1-dependent camalexin synthesis via PEPR-mediated signaling.

**Figure 3:**
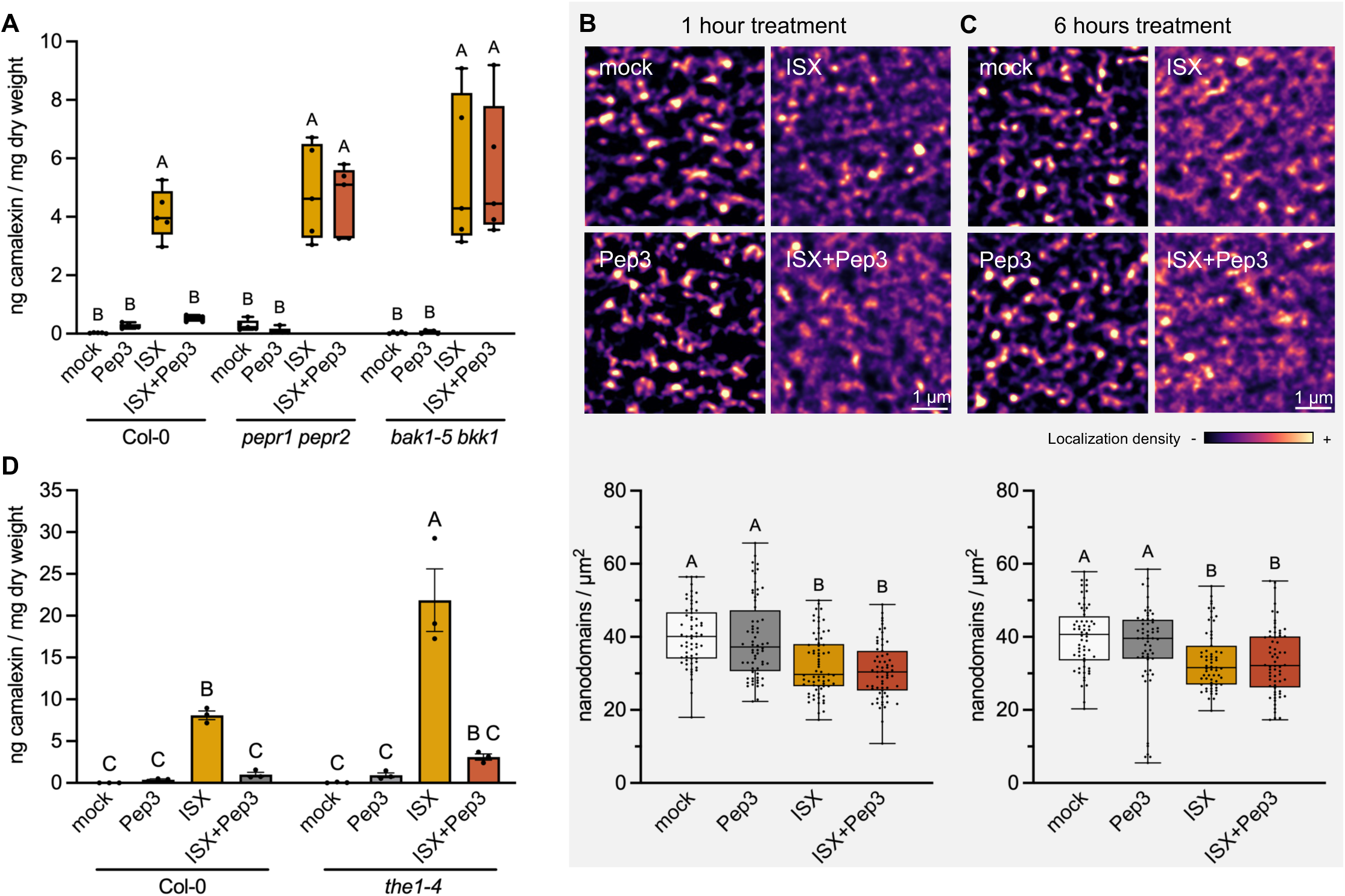
Pep3-PEPR signaling impairs induction of camalexin synthesis downstream of THE1. (A) Camalexin amount in Col-0, *pepr1 pepr2* and *bak1-5 bkk1* seedlings at 8 h after mock, Pep3, isoxaben (ISX) and ISX+Pep3 treatment (n=5). Different letters indicate statistically significant differences according to two-way ANOVA and Holm-Sidak’s multiple comparisons test (α = 0.05). (B,C) THE1-GFP nanodomain density was determined in seedling hypocotyls at (B) 1 h and (C) 6 h after treatment with mock, Pep3, ISX and ISX+Pep3. Representative images of enhanced super-resolution radial fluctuation (eSRRF) analysis are shown alongside the quantification of nanodomain density in 3 pooled experiments with 4-8 cells from 3 seedlings per experiment (n=58-69). Different letters indicate statistically significant differences according to Kruskal-Wallis and Dunn’s test (α = 0.05). (D) Camalexin amount in Col-0 and *the1-4* seedlings at 8 h after mock, Pep3, ISX and ISX+Pep3 treatment (n=3). Error bars indicate the SEM. Different letters indicate statistically significant differences according to two-way ANOVA and Holm-Sidak’s multiple comparisons test (α = 0.05).

Despite the recognized function of THE1 as a CWI sensor, little is known about its regulation upon perception of CWD. Accumulating reports link changes in the plasma membrane protein organization with the initiation of cell signaling (Jaillais et al., 2024). We combined enhanced super-resolution radial fluctuation (eSRRF) (Laine et al., 2023) with variable-angle total internal reflection microscopy (VA-TIRFM) imaging (Konopka and Bednarek, 2008) to analyze the plasma membrane organization of THE1-GFP upon CWD. We observed that a subpopulation of THE1-GFP was organized into nanodomains and that 1 hour and 6 hours of ISX treatment led to a decrease in the density of THE1-GFP nanodomains (Figure 3B,C). Congruently, the relative proportion of THE1-GFP localization organized in nanodomains and the image-wide spatial clustering index (iSCI) of THE1-GFP were reduced at both time points after ISX treatment (Figure S4). Confocal microscopy showed that THE1-GFP localization to the plasma membrane was unaffected by ISX treatment (Figure S5). Together, these data indicate that THE1-GFP is spatially redistributed within the plasma membrane upon CWD induced by ISX and may indicate that THE1-GFP nanodomains correspond to resting pools of the receptor that are released for function. To test whether the Pep3-dependent suppression of THE1-mediated camalexin synthesis could be linked to altered THE1 localization, we treated THE1-GFP seedlings with ISX, Pep3 and a combination of both. Neither the mean THE1-GFP fluorescence at the plasma membrane nor THE1 nanodomain organization were altered by Pep3 co-treatment, suggesting that negative regulation of ISX responses is exerted downstream of THE1 (Figure 3B,C, Figure S4, Figure S5). In good agreement with this conclusion, suppression of ISX-induced camalexin by Pep3 was also observed in the hypermorphic *the1-4* allele (Figure 3D).

### Pep3-dependent inhibition of CWI responses is independent from WRKY33 and MPK3/6

To investigate if Pep3 inhibition of CWI signaling downstream of THE1 involves transcriptional regulation of *WRKY33* or *PAD3*, we performed time course qRT-PCR analysis in Col-0, *the1-1*, *the1-4* and *pepr1 pepr2* mutants. Pep3 treatment led to upregulation of *WRKY33* after 2 hours, whereas *PAD3* expression was not induced at this time point (Figure S6). ISX caused the upregulation of *WRKY33* between 2 and 4 hours after treatment and of *PAD3* between 4 and 6 hours after treatment (Figure S6). Notably, ISX-induced *WRKY33* induction was neither significantly affected by THE1, nor by Pep3 co-treatment (Figure 4A). Instead, *PAD3* induction after ISX treatment was reduced in *the1-1,* increased in *the1-4* and suppressed by Pep3 co-treatment, showing that THE1 and Pep3 regulate *PAD3* without influencing *WRKY33* expression (Figure 4B).

**Figure 4:**
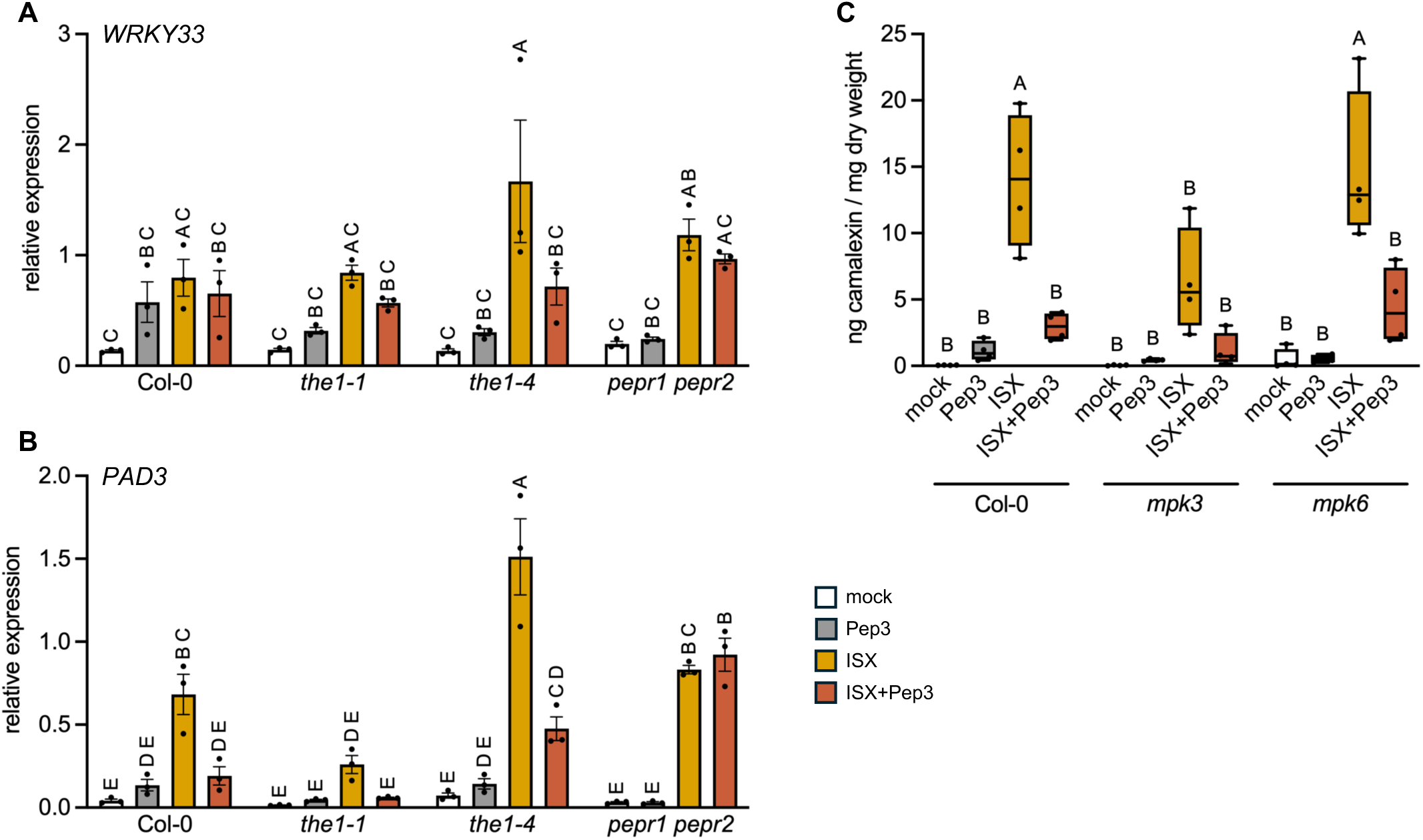
Pep3-dependent inhibition of CWI signaling does not involve WRKY33 and MPK3/6. Relative expression of (A) *WRKY33* and (B) *PAD3* was determined in Col-0, *the1-1*, *the1-4* and *pepr1 pepr2* at 6 h after treatment with mock, Pep3, isoxaben (ISX) and ISX+Pep3 via quantitative RT-PCR. Values are relative to *ACT2* expression and error bars represent the SEM (n=3). (C) Camalexin amount in Col-0, *mpk3* and *mpk6* seedlings at 8 h after mock, Pep3, ISX and ISX+Pep3 treatment (n=4). Different letters indicate statistically significant differences according to two-way ANOVA and Holm-Sidak’s multiple comparisons test (α = 0.05).

We previously reported that the MAP kinase kinase kinase (MAPKKK) family proteins Arabidopsis NPK1-related Protein kinase 1 (ANP1) and ANP3 regulate ISX-dependent responses, indicating that MAP kinase signaling plays a role in CWI signaling (Gigli Bisceglia et al., 2018). Camalexin accumulation after pathogen infection involves phosphorylation of WRKY33 by the MAP kinases MPK3 and MPK6, both of which are phosphorylated by Pep3-triggered signaling (Mao et al., 2011; Bartels et al., 2013). To test if MPK3/6 are involved in ISX-induced camalexin synthesis and/or in Pep3-dependent inhibition of the same, we analyzed camalexin levels in Col-0, *mpk3* and *mpk6* seedlings after treatment with mock, Pep3, ISX and ISX+Pep3. MPK3, but not MPK6, was required for ISX-induced camalexin accumulation (Figure 4C). We were however unable to detect any changes in MPK3/6 phosphorylation after ISX treatment for 10 min, when CESAs are internalized (Paredez et al., 2006), as well as after 4 and 6 hours, when transcriptional responses become more pronounced and phytohormones and camalexin start to accumulate (Denness et al., 2011; Engelsdorf et al., 2018; Zhai et al., 2024) (Figure S7). While Pep3 single or combined treatments induced MPK3 and MPK6 phosphorylation (Figure S7), Pep3-dependent suppression of ISX-induced camalexin accumulation was not affected in *mpk3* and *mpk6* mutants, suggesting that MPK3/6 signaling is not required for Pep3-triggered inhibition of CWI responses (Figure 4C).

Taken together, our data indicate that Pep3-PEPR signaling inhibits ISX-induced production of camalexin downstream of THE1 and independent from MPK3/6 and WRKY33.

### MYB and WRKY transcription factors are regulated by THE1 and Pep3 signaling

To identify candidate genes that mediate Pep3 inhibition of CWI signaling, we performed RNA-Sequencing of Col-0 seedlings treated with mock, Pep3, ISX and ISX+Pep3 for 6 hours, when *PAD3* expression was suppressed by the co-treatment (Figure 4B). Principal component analysis showed that biological replicates grouped according to treatments and indicated that the largest share of the transcriptional changes in the data sets could be explained by ISX treatment (Figure S8A). Hierarchical cluster analysis supported this conclusion by grouping ISX and ISX+Pep3 treatments together but also revealed groups of genes with opposite expression patterns between both treatments (Figure S8B). Analysis of differential gene expression identified a total of 543 genes significantly downregulated (α=0.05, fold change > 2) after Pep3 (89 genes), ISX (330) and ISX+Pep3 (313) treatment compared to mock control (Figure 5A). Instead, 1513 genes were upregulated (α=0.05, fold change > 2) after treatment with Pep3 (416 genes), ISX (1248) and ISX+Pep3 (1104) (Figure 5B). The transcription factors *MYB47*, *MYB95*, *MYB113* and *WRKY67* were significantly upregulated by ISX, but not after combined ISX+Pep3 treatment, making them candidates for mediating the negative effect of Pep3 on CWI signaling. Quantitative RT-PCR showed that all four candidates were induced in a THE1-dependent manner, with significantly increased expression in ISX-treated *the1-4* and reduced expression in *the1-1* compared to the wild type (Figure 5C-F). *MYB47*, *MYB95* and *MYB113* expression were mildly reduced by loss of *WRKY33*, whereas *WRKY67* expression was increased in *wrky33*, *the1-1 wrky33* and *the1-4 wrky33* compared to Col-0 and *the1* single mutants, respectively (Figure 5C-F). To investigate if MYB47 and the closely related MYB95 (Millard et al., 2019) are involved in the regulation of ISX-dependent camalexin accumulation by Pep3, we subjected *myb47* single and *myb47/95* double mutants to mock, Pep3, ISX and combined ISX and Pep3 treatments. We detected significantly reduced camalexin levels after ISX treatment in both lines, indicating that MYB47 is required for induction of camalexin biosynthesis, with no obvious contribution of MYB95 (Figure 5G,H). Congruently, the ISX-induced expression of *PAD3* was significantly reduced in *myb47* and *myb47/95* seedlings, whereas the induction of *CYP71A13* was only reduced in *myb47/95* seedlings (Figure 5I,J). Two independent mutant lines for *MYB113* did not cause significant changes in camalexin accumulation compared to WT (Figure S9). We also obtained three independent T-DNA insertion lines for *WRKY67*, of which only SALK_073815 (now termed *wrky67-1*) displayed reduced gene expression after ISX treatment, but no change in ISX-induced camalexin accumulation (Figure S9).

**Figure 5:**
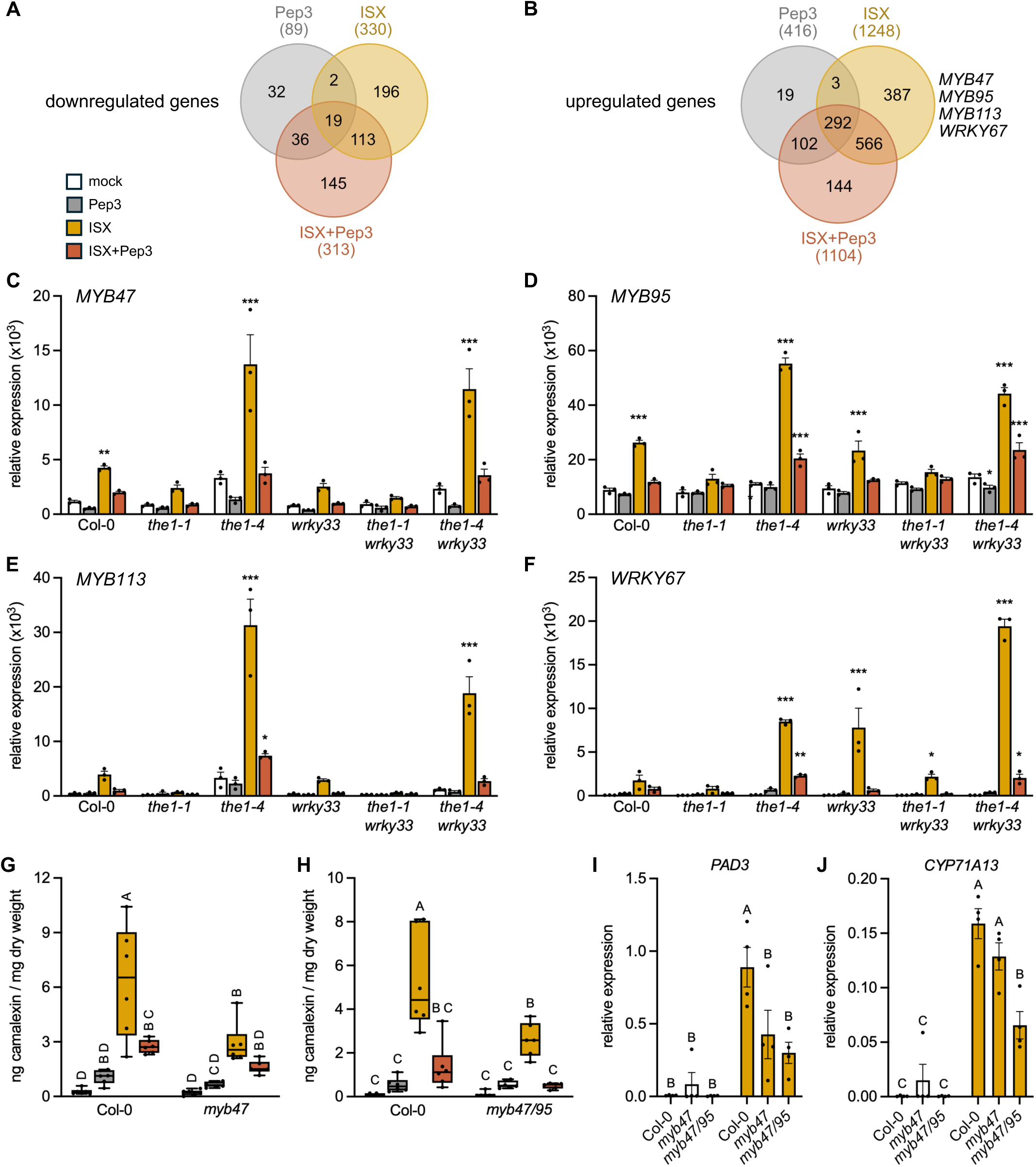
Identification of Pep3-regulated transcription factors involved in ISX-induced camalexin accumulation. (A, B) RNA-Sequencing of Col-0 seedlings identifies genes (A) upregulated and (B) downregulated at 6 h after treatment with mock, Pep3, isoxaben (ISX) and ISX+Pep3 (α = 0.05, fold change >2). (C-F) Relative expression of (B) *MYB47*, (C) *MYB95,* (D) *MYB113* and (E) *WRKY67* was determined in Col-0, *the1-1*, *the1-4, wrky33, the1-1 wrky33* and *the1-4 wrky33* at 6 h after treatment with mock, Pep3, ISX and ISX+Pep3 via quantitative RT-PCR. Values are relative to *ACT2* expression and error bars represent the SEM (n=3). Asterisks indicate statistically significant differences to the respective mock treatments according to two-way ANOVA and Holm-Sidak’s multiple comparisons test (*p < 0.05, **p < 0.01, ***p < 0.001). (G,H) Camalexin amount in (G) Col-0 and *myb47,* and in (H) Col-0 and *myb47/95* seedlings at 8 h after mock, Pep3, ISX and ISX+Pep3 treatment (n=6). (I,J) Relative expression of (I) *PAD3* and (J) *CYP71A13* was determined in Col-0, *myb47* and *myb47/95* seedlings at 8 h after treatment with mock and ISX via quantitative RT-PCR. Values in are relative to *ACT2* expression and error bars represent the SEM (n=4). Different letters in (G-J) indicate statistically significant differences according to two-way ANOVA and Holm-Sidak’s multiple comparisons test (α = 0.05).

Taken together, our RNA-Seq analysis identified four transcription factors that were induced in a THE1-dependent manner and suppressed by Pep3 co-treatment, of which *MYB47* and *MYB95* were required for the regulation of camalexin biosynthesis after CWD induction by ISX treatment.

### MYB47/95 mediate THE1-induced and Pep3-sensitive JA signaling

Previous research connected JA to camalexin biosynthesis induction and demonstrated that JA accumulation in response to ISX treatment is sensitive to Pep signaling (Denness et al., 2011; Engelsdorf et al., 2018; Zhou et al., 2022). This overlap prompted us to investigate the role of JA in ISX-induced camalexin production and its Pep3-dependent negative regulation. Our transcriptomics data showed an increased expression of genes involved in JA biosynthesis, signaling and catabolism after 6 hours of ISX treatment, reduced expression after 6 hours Pep3 treatment, and an attenuated expression profile after combined ISX+Pep3 treatment compared to ISX treatment alone (Figure 6A). JA-deficient *aos* mutants did not accumulate camalexin after ISX treatment, supporting an essential role of JA (Figure 6B). Since JA induction after ISX treatment is attenuated by Pep3 (Engelsdorf et al., 2018), we hypothesized that exogenous application of JA might abrogate Pep3-dependent suppression of camalexin production. Indeed, triple treatment with ISX, Pep3 and JA strongly induced camalexin accumulation, demonstrating that JA application is sufficient to override the negative influence of Pep3 (Figure 6C).

**Figure 6:**
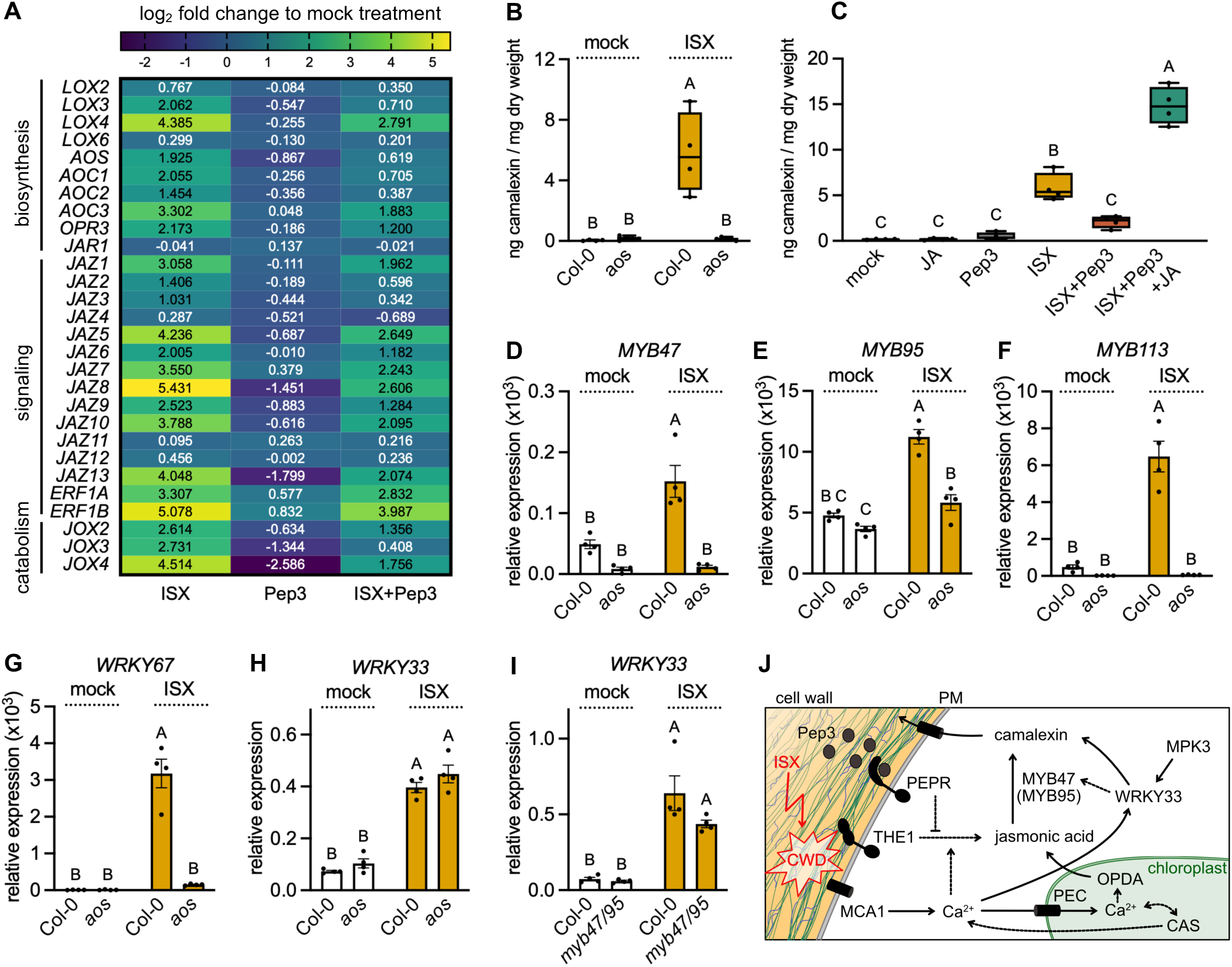
ISX-induced and Pep3-sensitive camalexin accumulation depends on jasmonic acid. (A) Transcripts involved in jasmonic acid (JA) biosynthesis, signaling and catabolism were analyzed via RNA-Sequencing in Col-0 seedlings treated with mock, Pep3, isoxaben (ISX) and ISX+Pep3 for 6 h. Relative expression after ISX, Pep3 and ISX+Pep3 compared to mock treatment is listed as log_2_ fold change and visualized as heat map indicating downregulation (dark blue) and upregulation (yellow) of transcripts. (B, C) Camalexin amount in (B) Col-0 and *aos* seedlings at 8 h after mock and ISX treatment (n=4-5) and (C) Col-0 seedlings at 8 h after mock, JA, Pep3, ISX, ISX+Pep3 and ISX+Pep3+JA treatment (n=4). (D-G) Relative expression of (D) *MYB47*, (E) *MYB95,* (F) *MYB113* and (G) *WRKY67* was determined in Col-0 and *aos* seedlings at 6 h after treatment with mock and ISX via quantitative RT-PCR. (H,I) Relative expression of *WRKY33* was determined in (H) Col-0 and *aos* and (I) Col-0 and *myb47/95* seedlings at 6 h after treatment with mock and ISX via quantitative RT-PCR. Values in (D-I) are relative to *ACT2* expression and error bars represent the SEM (n=4). Different letters indicate statistically significant differences according to (C) one-way ANOVA or (B,D-I) two-way ANOVA and Holm-Sidak’s multiple comparisons test (α = 0.05). (J) Model depicting the induction of camalexin synthesis after perception of cell wall damage (CWD) via THE1, JA and MYB47/95. Interactions with Pep3/PEPR-, WRKY33-, and Ca^2+^ signaling are proposed based on data presented in this manuscript and references discussed in the main text.

Next, we investigated if ISX-induced expression of *MYB47*, *MYB95*, *MYB113* and *WRKY67* depends on JA biosynthesis. Quantitative RT-PCR showed that ISX-dependent induction of *MYB47*, *MYB113* and *WRKY67* was abolished and induction of *MYB95* attenuated in JA-deficient *aos* mutants (Figure 6D-G). ISX-induced expression of *WRKY33* was independent of JA and MYB47/95 (Figure 6H,I). Together, our data support a model in which THE1-dependent CWI surveillance induces camalexin via JA and MYB47/95, whereas an independent pathway induces *WRKY33* upon CWD (Figure 6J).

## Discussion

In this study, we investigated contributions of THE1-dependent CWI signaling to the induction of the phytoalexin camalexin and its interaction with Pep3 signaling. We show that CWI impairment caused by cellulose biosynthesis inhibition, enzymatic cell wall degradation or fungal infection induces camalexin accumulation. In line with the previously described role of WRKY33 as central regulator of camalexin biosynthesis (Mao et al., 2011), increased CWI signaling in the hypermorphic *the1-4* mutant required WRKY33 for camalexin induction. WRKY33 expression was induced by ISX treatment, but was neither dependent on THE1, nor affected by co-treatment with Pep3. This is in contrast with ISX-induced expression of PAD3, catalyzing the final step of camalexin synthesis, which strongly depended on THE1 and was suppressed by co-treatment with Pep3. Phosphorylation by MPK3/6 and CALCIUM-DEPENDENT PROTEIN KINASEs (CPK5/6) controls WRKY33 DNA binding, autoactivation, and transactivation of key camalexin biosynthesis genes (Zhou et al., 2020). Our analysis showed that MPK3 controls ISX-induced camalexin accumulation, while MPK6 was not required. MPK3 phosphorylation was not detectable at 10 minutes, 4 hours and 6 hours after ISX treatment, indicating that (i) MPK phosphorylation after ISX treatment is not a fast response, and (ii) that it may represent a transient and/or dynamic response that is difficult to catch.

The importance of calcium signaling for ISX-induced CWD responses has long been recognized (Denness et al., 2011), but the exact mode of action remains elusive. Current evidence suggests that MCA1 might represent a stretch-activated Ca^2+^ channel required for THE1-dependent CWI signaling (Nakagawa et al., 2007; Engelsdorf et al., 2018). Delayed inhibitor treatment assays with LaCl_3_ showed that calcium signaling affects responses to ISX treatment that occur in the scale of hours, when CWD causes mechanical damage and microscopically visible phenotypes (Denness et al., 2011; Engelsdorf et al., 2018; Bacete et al., 2022). Congruently, LaCl_3_ treatment abolished ISX-induced camalexin accumulation. Loss of MCA1 halved camalexin levels compared to wild type, indicating that it may be partially responsible for Ca^2+^ influx after CWD. Similarly, CAS, a chloroplast-localized positive regulator of Ca^2+^ responses in the cytosol and the chloroplast stroma (Nomura et al., 2008; Weinl et al., 2008; Nomura et al., 2012), contributed to camalexin accumulation in response to ISX. Loss of chloroplast inner envelope ion channels PEC1/2 impairs stromal Ca^2+^ transients but increases cytosolic Ca^2+^ transients, suggesting that Ca^2+^ is partially excluded from plastids (Völkner et al., 2021). Our data indicate that increased cytosolic Ca^2+^ in *pec1/2* triggered ISX-induced camalexin accumulation. While CWI maintenance during high salinity requires the *Cr*RLK1L FER to induce late Ca^2+^ transients (Feng et al., 2018), *fer* mutants exhibit constitutively increased CWD responses that cannot be alleviated by loss of THE1 (Engelsdorf et al., 2018; Gonneau et al., 2018), indicating that different Ca^2+^ signatures might be involved in CWI maintenance and CWD responses. We provide first hints that CWD increases responsiveness to cell wall-related Ca^2+^ elicitation as ISX treated seedlings responded stronger to high salinity and cold treatments. It will be interesting to investigate in the future if CWI sensors contribute to this response.

In recent years, plasma membrane localized proteins have been recognized to be dynamically organized in specific nanodomains according to developmental programs and stress responses (Bücherl et al., 2017; Hdedeh et al., 2025). Protoplast cell wall regeneration assays indicated that the cell wall influences lateral mobility of plasma membrane proteins and chemical treatments affecting cellulose biosynthesis or pectin methylation caused changes in protein nanodomain size and dynamics (Martinière et al., 2012; McKenna et al., 2019). These changes might influence signaling, as the nanodomain organization of RLKs is influenced by FER, cell wall localized LRR-EXTENSIN (LRX) and their ligand RALF23 (Gronnier et al., 2022; Liu et al., 2024). In line with its central function in CWI signaling upon CWD, THE1 was spatially reorganized at the plasma membrane after one hour of ISX treatment, showing that THE1 reorganization precedes accumulation of phytohormones and camalexin. This observation suggests that THE1 may be released from nanodomains for function. Similarly, during viral infection, the solicitation of REMORIN proteins correlates with their spatial redistribution and an overall increase in their lateral diffusion (Perraki et al., 2018; Jolivet et al., 2025). Because ligand-induced signaling can influence nanodomain organization of other receptor kinases (Gronnier et al., 2022), we considered the possibility that Pep3 signaling may influence THE1 nanodomain organization. However, we observed no effects of Pep3 treatments on THE1-GFP nanodomains under mock or ISX conditions, indicating interaction of CWI and Pep3 pathways downstream of THE1. Consistently, we observed that Pep3 inhibited ISX-induced camalexin accumulation in the hypermorphic *the1-4* allele.

To identify potential downstream regulators, we performed transcriptomics and discovered four candidate transcription factors that were induced by ISX treatment in a THE1-dependent manner and suppressed by Pep3 co-treatment, indicating that they are downstream of a Pep3-THE1 pathway interaction. Mutant analysis revealed that *MYB47* and *MYB95* are required for the induction of camalexin biosynthesis upon ISX treatment, extending the network of transcription factors known to be involved in the regulation of the camalexin pathway. MYB34, MYB51 and MYB122 regulate *CYP79B2* and *CYP79B3*, thereby affecting the conversion of tryptophan to IAOx and determining precursor availability for camalexin synthesis (Frerigmann et al., 2015). Instead, ANAC042, ERF1B and ERF72 are required for transcriptional induction of *CYP71A13* and *PAD3*, directly regulating the camalexin biosynthesis pathway (Saga et al., 2012; Li et al., 2021; Zhou et al., 2022; Monsalvo et al., 2025). Of note, ERF1B cooperates with WRKY33 to induce camalexin pathway genes and is induced by JA (Zhou et al., 2022). These properties are reminiscent of THE1 signaling in response to CWD, which induces the JA pathway and acts in parallel to WRKY33 in camalexin synthesis. Indeed, JA was required for ISX-induced camalexin accumulation as well as for induction of *MYB47*, *MYB95*, *MYB113* and *WRKY67.* In Arabidopsis roots, cellulose defects cause cortex cell bulging and compression of inner tissues stimulating JA signaling (Mielke et al., 2021). The same mechanism might be responsible for JA induction after ISX treatment, where THE1 is required for CWD detection. *MYB47* induction was previously detected in drought treated Arabidopsis plants, where it was associated with increased JA signaling and drought stress survival (Marquis et al., 2022). Since drought stress impairs CWI (Novaković et al., 2018), it would be interesting to test if *MYB47* expression upon drought is induced by THE1-dependent CWI signaling. Recently, Cao et al. (2025) reported that *MYB47* is induced by JA and increases JA catabolism in leaves, thereby delaying leaf senescence. Bizan et al. (2025) showed that *MYB47* and *MYB95* are controlled by master transcription factors of JA signaling, MYC2/3/4, and regulate a specific subset of JA-responsive genes related to ER-body formation and defense against fungal and insect infection. JA treatment alone was not sufficient to induce camalexin synthesis, and the available data indicates that at least two independent pathways need to be activated upon CWD to initiate this defense response (Zhou et al., 2022). Consistently, negative regulation of JA synthesis by Pep3 treatment was sufficient to suppress the camalexin pathway whereas JA application overrode the suppressive effect of Pep3 on ISX-induced camalexin accumulation.

Taken together, perception of CWD involves changes in THE1 nanodomain organization at the plasma membrane and regulation of Ca^2+^ signaling via MCA1, CAS and PEC (Figure 6J). THE1-induced JA signaling and THE1-independent WRKY33 are both induced by CWD and required for camalexin accumulation. Negative regulation of this cell wall-mediated defense response by PEPR-mediated perception of Pep3 is placed upstream of JA. We identified *MYB47*, *MYB95*, *MYB113* and *WRKY67* as ISX-induced, THE1- and JA-dependent transcription factors, of which *MYB47* and *MYB95* were required for the regulation of ISX-induced camalexin biosynthesis. We propose that the induction of camalexin production by CWD may contribute to defense at fungal penetration sites. In turn, current evidence suggests that Peps are released upon plasma membrane rupture allowing their binding to PEPRs of neighboring cells (Hander et al., 2019; Shen et al., 2019). Pep3 detection may therefore signify damage on the tissue level, leading to suppression of local cell wall-mediated defense at infection sites.

## Materials and Methods

### Plant materials and growth conditions

*Arabidopsis thaliana* genotypes used in this study are listed in Supplemental Table S1. Accession Columbia-0 (Col-0) was used as wild type in all experiments. All Arabidopsis mutants used in this study were either obtained from the Nottingham Arabidopsis Stock Centre (http://arabidopsis.info/), from laboratories where data on them has been published, or have been crossed from those.

For growth in sterile liquid culture, seeds were surface sterilized for 10 minutes with 70 % (v/v). The ethanol was subsequently replaced with a sterilizing solution of 2% sodium hypochlorite with 0.1 % Triton X-100, treated for additional 3 minutes and finally rinsed four times with sterile water. Seeds were then stratified at 4°C for 3 days before transfer to half-strength Murashige and Skoog liquid medium [Murashige and Skoog Basal Medium (2.1 g/liter), MES salt (0.5 g/liter), and 1 % sucrose, pH 5.7]. Unless stated otherwise, seedling cultures were established in a 24-well plate with 6 seeds per well. Seedlings were grown under a 16-hour light (22°C) / 8-hour dark (20°C) cycle and a photon flux density of 110 μmol m^−2^ s^−1^.

For growth on soil, Arabidopsis seeds were placed on a mixture of 80 % Fruhstorfer Type P (Hawita, Germany) and 20 % Liadrain clay (Liapor, Germany), stratified for 3 days and grown at a photon flux density of 110 μmol m^−2^ s^−1^ under the light cycles indicated in the figure legends.

### Plant liquid culture treatments

Unless stated otherwise, 10-day-old seedlings were subjected to the following treatments in fresh liquid growth medium at the indicated final concentrations: 600 nM isoxaben (ISX; Supelco 36138), 10 nM *At*Pep3, 600 nM ISX + 10 nM *At*Pep3 (combined treatment of ISX and AtPep3), 10 mM LaCl_3_, 5 μM jasmonic acid (JA; Cayman Chemical 88300), 600 nM ISX + 10 nM *At*Pep3 + 5 μM JA (combined treatment of ISX, *At*Pep3, and JA). Equal amounts of DMSO were present in all treatments, including mock treatments. Treatment with 0.03 % (w/v) Driselase (Dri; Sigma-Aldrich D8037) and inactivated (bDri) was performed according to Engelsdorf et al. (2018). After treatment, seedlings were briefly rinsed, patted dry, snap-frozen in liquid nitrogen and stored at -70°C until further analysis. *At*Pep3 (EIKARGKNKTKPTPSSGKGGKHN) was custom synthesized and supplied by EZBiolab Inc. (Indiana, USA).

### *Colletotrichum higginsianum* infection assays

*Colletotrichum higginsianum* isolate MAFF 305635 (Ministry of Forestry and Fisheries, Japan) was used for all infection experiments. Conidia were plated on oatmeal agar (5 % [w/v] shredded oatmeal, 1.2 % [w/v] agar) and incubated for 7 days in a 16 h light (22°C) / 8 h dark (20°C) cycle at a photon flux density of 110 µmol m^-2^ s^-1^. Immediately prior to infection, conidia were harvested by rinsing plates with deionized water and the titer was adjusted to 2 · 10^6^ conidia ml^-1^. Spray infection of leaf rosettes was performed at the end of the light phase and high humidity maintained until 2.5 days post infection to support fungal penetration as described (Voll et al., 2012). Fungal entry rates were determined by microscopic examination after staining fungal structures with trypan blue as described in Engelsdorf et al. (2017) and quantification of the relative fungal DNA content in infected leaves was performed by qPCR analysis of *ChTrpC* as described in Engelsdorf et al. (2013).

### Quantitative reverse transcription polymerase chain reaction (qRT-PCR)

Total RNA was isolated using a NucleoSpin Plant and Fungi RNA Mini Kit (Macherey-Nagel). One microgram of total RNA was treated with RNase-Free DNase (Thermo Scientific) and used for complementary DNA (cDNA) synthesis with a RevertUP II Reverse Transcriptase Kit (Biotechrabbit). Quantitative reverse transcription polymerase chain reaction (qRT-PCR) was performed using Blue S’Green (Biozym) SYBR green mix and a Bio-Rad CFX Connect RT-PCR system. Primers are listed in Supplemental Table S2. *ACT2* was used a reference in all experiments.

### RNA-Sequencing

Total RNA was isolated using a NucleoSpin Plant and Fungi RNA Mini Kit (Macherey-Nagel) and shipped to Novogene Europe (Cambridge, UK) for further processing. RNA integrity was assessed using the RNA Nano 6000 Assay Kit of the Bioanalyzer 2100 system (Agilent Technologies, CA, USA). Total RNA was used as input material for the RNA sample preparations. Briefly, mRNA was purified from total RNA using poly-T oligo-attached magnetic beads. Fragmentation was carried out using divalent cations under elevated temperature in First Strand Synthesis Reaction Buffer (5X). First strand cDNA was synthesized using random hexamer primer and M-MuLV Reverse Transcriptase (RNase H-). Second strand cDNA synthesis was subsequently performed using DNA Polymerase I and RNase H. Remaining overhangs were converted into blunt ends via exonuclease/polymerase activities. After adenylation of 3’ ends of DNA fragments, adaptors with hairpin loop structure were ligated to prepare for hybridization. To select cDNA fragments of preferentially 370∼420 bp in length, the library fragments were purified with AMPure XP system (Beckman Coulter, Beverly, USA). PCR was performed with Phusion High-Fidelity DNA polymerase using universal and index PCR primers, PCR products were purified (AMPure XP system), and library quality was assessed on the Agilent Bioanalyzer 2100 system. The clustering of the index-coded samples was performed on a cBot Cluster Generation System using TruSeq PE Cluster Kit v3-cBot-HS (Illumina) according to the manufacturer’s instructions. After cluster generation, the library preparations were sequenced on an Illumina Novaseq platform and 150 bp paired-end reads were generated. Raw data (raw reads) of fastq format were firstly processed through in-house perl scripts. In this step, clean data (clean reads) were obtained by removing reads containing adapter, reads containing poly-N and low-quality reads from raw data. All the downstream analyses were based on the clean data with high quality. Index of the TAIR10 reference genome was built using Hisat2 v2.0.5 and paired-end clean reads were aligned to the reference genome using Hisat2 v2.0.5. Differential expression analysis was performed using the DESeq2 R package (1.20.0). The resulting p-values were adjusted using the Benjamini and Hochberg’s approach for controlling the false discovery rate. Genes with an adjusted p-value <=0.05 found by DESeq2 and a fold change > 2 were assigned as differentially expressed.

### Camalexin quantification

To analyze camalexin content in seedlings grown in liquid culture, whole seedlings were freeze-dried for 17 hours using the CHRIST ALPHA 2-4 system (Martin Christ, Osterode am Harz, Germany) and their dry weight recorded. For *Colletotrichum higginsianum*-infected leaf material, camalexin was directly extracted from sampled leaf discs as described (Nawrath and Métraux, 1999; Engelsdorf et al., 2013). Crude phytochemicals were extracted from the samples in 600 µl 70 % methanol supplemented with 50 ng ortho-anisic acid (oANI) as internal standard at 65°C for 1 hour. Crude extraction was repeated on the same samples with 90 % methanol at 65°C for 1 hour. The two extractions were combined, and a rotatory vacuum concentrator (RVC 2-25 CDplus, Martin Christ) was used to evaporate methanol. Proteins and other biomolecules were precipitated from the extract with 500 μl of 5 % trichloroacetic acid. After centrifugation, the supernatant was collected into a fresh reaction tube and partitioned twice against 600 μl of cyclohexane/ethyl acetate (1:1 (v/v)) solution with vigorous shaking and centrifugation at 3,500*g* for 10 min. The combined organic (upper) phases were evaporated in the vacuum concentrator and resuspended in 80 % 25 mM potassium dihydrogen phosphate buffer (25 mM KH_2_PO_4_, pH 2.6) / 20% acetonitrile (ACN). Camalexin content in the extract was separated on an InfinityLab Poroshell 120 EC-C18 (3.0 x 150 mm 2.7-Micron, Agilent Technologies) column and analyzed using an UHPLC system (1260 Infinity II, Agilent Technologies, Santa Clara, California, USA). The separation was achieved using gradient elution of a mobile phase consisting of (A) 25 mM KH_2_PO_4_, pH 2.6, and (B) 99 % ACN in water at a flow rate of 1 ml/min: 0 – 2.5 min: 20 % B; 2.5 – 6 min: 20 % to 80 % B; 6 – 7 min: 80 % B; 7 – 8 min: 80 % to 20 % B; 8–10 min: 20 % B. The G712B fluorescence detector (Agilent Technologies) was used to detect oANI (excitation 305 nm, emission 365 nm) and camalexin (excitation 305 nm, emission 370 nm). A standard dilution series of oANI and camalexin in the range of 0.02 to 100 ng was employed for quantification. OpenLAb CDS ChemStation software C01.10 (Agilent Technologies) was used to operate the HPLC system and to analyze the data.

### MAP kinase phosphorylation assay

Seedlings were grown in half-strength Murashige and Skoog (MS) liquid medium [Murashige and Skoog Basal Medium (2.1 g/L), MES (0.5 g/L), and 1 % sucrose, pH 5.7] in 12-well plates. On the ninth day, the medium was replaced with fresh medium, and on the following day seedlings were treated with either Mock (DMSO), 600 nM ISX, 10 nM *At*Pep3, or a combined treatment of 600 nM ISX + 10 nM *At*Pep3 for 10 min, 4 hours, or 6 hours. Protein extraction was performed as described in Gigli-Bisceglia et al. (2022). For protein extraction, the following buffer was used: 50 mM Tris-HCl (pH 7.5), 200 mM NaCl, 1 mM EDTA, 10 % (v/v) glycerol, 0.1 % (v/v) Tween 20, 1 mM PMSF, 1 mM DTT, 1× protease inhibitor cocktail (Sigma-Aldrich, P9599), and 1× phosphatase inhibitor cocktail (Sigma-Aldrich, P2850). Samples were incubated on ice for 15 min and centrifuged at 13,000 g for 15 min at 4°C. The resulting supernatants were used for western blot analysis. Protein concentration was determined using the Bradford assay (Bio-Rad). Samples were mixed with 4× loading buffer [240 mM Tris-HCl (pH 6.8), 8 % (w/v) SDS, 40 % glycerol, 5 % β-mercaptoethanol, 0.04 % (w/v) Bromophenol Blue] and heated at 95°C for 2 min. Protein samples (35 µg total protein per lane) were separated on 10 % SDS-PAGE gels and transferred to nitrocellulose membranes using the Trans-Blot Turbo Transfer System (Bio-Rad) at 25 V, 1.3 A for 7 min. Membranes were blocked in Tris-buffered saline with 0.1 % Tween 20 (TBST) containing 5 % (w/v) bovine serum albumin (BSA; Sigma-Aldrich, A3608) for 1 h, then incubated overnight with the primary phospho-p44/42 MAPK (Erk1/2) (Thr202/Tyr204) antibody (1:2000, Cell Signaling Technology, 9101). After three washes with TBST, membranes were incubated with the secondary HRP-linked anti-rabbit IgG antibody (1:6000, Cell Signaling Technology, 7074) for 2 hours. Following three additional washes with TBST, chemiluminescence was detected using Clarity Western ECL Substrate (Bio-Rad, 1705061) for 2 min and visualized with the ChemiDoc imaging system (Bio-Rad). For equal loading analysis, membranes were stripped using stripping solution (100 mM Tris-HCl, pH 6.8, 2 % SDS, 100 mM β-mercaptoethanol) for 1 hour at 60°C, re-blocked as described above, and probed with the β-Actin (C4) antibody (1:2000, Santa Cruz Biotechnology, sc-47778) for 1 hour. Following three washes (TBST), chemiluminescence detection was performed as described previously.

### Ca^2+^ assays

To monitor cytosolic Ca²⁺ transients, we employed Arabidopsis reporter lines expressing cytosolic-targeted aequorin in Col-0 (pMAQ2; Knight et al., 1991) and *pec1-1 pec2-1* background (unpublished). The measurements were carried out as described before by Tanaka et al. (2013), with minor alterations. Briefly, seedlings (5 days old) grown sterilely on half-strength Murashige and Skoog (½ MS) medium were transferred into white 96-well plates, each well containing 50 µL of coelenterazine buffer (10 µM coelenterazine, 1.4 mM CaCl₂, 20 mM KCl, 5 mM MES-KOH, pH 5.7). For aequorin reconstitution, seedlings were kept in the dark at 22 °C overnight. For priming experiments, ISX was added at a final concentration of 600 nM 4.5 hours prior to the measurement. Cytosolic Ca^2+^ transients were elicited with 200 mM NaCl or ice-cold water. Luminescence was recorded using a TECAN Spark plate reader (Tecan, Männedorf, Switzerland), with an integration time of 500 ms. To determine total aequorin content, seedlings were discharged using 10% ethanol (v/v) and 1 M CaCl₂. Calcium levels were calculated from luminescence signals as described by (Knight et al., 1996).

### Confocal microscopy

Five-day old Arabidopsis seedlings grown on solid ½ MS, 1% Sucrose, 0.8 % Agar, pH 5.8 plates under a 16-hour photoperiod were incubated in ½ MS pH 5.8 liquid solutions containing 600 nM isoxaben, 10 nM *At*Pep3, a combination of both or corresponding mock controls for the indicated durations. Seedlings were mounted in the same solution they were treated in and imaged using a Leica SP8 confocal microscope equipped with a HC PL 10x/0.3 objective, a white light laser and HyD detectors. GFP was excited using 488 nm wavelength and fluorescence emission was detected between 495 nm and 535 nm. Acquisition parameters were kept constant during experiments. Fluorescence intensity was measured using Fiji (Schindelin et al., 2012).

### Variable angle-total internal reflection microscopy (VA-TIRFM)

Arabidopsis seedlings were grown and treated as described for confocal microscopy. Five-day old seedlings were mounted in the same solution they were treated in between two coverslips and imaged using a custom-build microscope platform for variable angle total internal reflection microscopy (VA-TIRFM) (Rohr et al., 2024), equipped with an 100x objective NA 1.49 (Zeiss, 421190-9800-000), a polychromatic modulator (AOTF, AA OPTO-ELECTRONIC) and a sCMOS camera (Hamamatsu Photonics, ORCA-Flash4.0 V2). To ensure that the target laser power value always corresponds to the actual irradiance in the sample plane, the power of each laser line was calibrated before each experiment using a PM100D detector (Thorlabs) as previously described (von Arx et al., 2024). Samples were excited with 488 nm diode laser at 200 µW and 20 seconds stream acquisitions were recorded at 5 Hz for each cell. Acquisition parameters were kept constant during experiments.

### Enhanced-super resolution radial fluctuation analyses (eSRRF)

Super-resolved image reconstructions of THE1-GFP plasma membrane organization were computed using eSRRF (Laine et al., 2023) in Fiji (Schindelin et al., 2012). Stream VA-TIRFM acquisitions were processed using the following parameters that were defined by iterative parameter sweep (Laine et al., 2023): magnification = 6, radius = 3, sensitivity = 3 and frames = 100. The average (AVG) reconstructions of THE1-GFP plasma membrane organization were segmented and analysed using Nanonet, an automated custom-made analysis pipeline (https://github.com/NanoSignalingLab/NanoNet), providing nanodomain density and image wide spatial clustering index (iSCI) based on predicted localization density.

### Statistical analysis

All statistical analysis were performed with GraphPad Prism (Version 10.5.0). Specific information on statistical methods, sample size, significance level and p-values are provided in the respective figure legends.

## Acknowledgements

This work was funded by the Deutsche Forschungsgemeinschaft (DFG, German Research Foundation) grants EN 1071/3-1 (to TE), A09-SFB1101 (to JG) and B01-TRR356 (to JG). H-HK and SM were supported by DFG SFB-TR 175, project B09 (INST 86/2119-2). NGB was supported by the Dutch Research Council (NWO) under grant numbers OCENW.M.24.079 and OCENW.XS23.1.050.

We are grateful to Debora Gasperini (IPB Halle) for providing *aos* seeds; Stefan Weinl and Jörg Kudla (Universität Münster) for *cas-1* seeds; Erich Glawischnig (TU Munich) for *cyp71a12/a13* seeds; and Kay Schneitz (TU Munich) for *pTHE1::THE1-GFP* seeds. Live cell imaging was performed at the ZMBP microscopy facility of the University of Tübingen and at the Center for Advanced Light Microscopy (CALM) of the TUM School of Life Sciences. We thank Michelle von Arx (TU Munich) for providing access to the NanoNet software prior publication and acknowledge Christiane Rohrbach and Nadja Braun (both Philipps-Universität Marburg) for excellent technical assistance.

## Author contributions

Conceptualization and project administration: TE; Investigation: RNM, LF, JD, DB, SM, AR, NGB, TE; Funding acquisition: H-HK, JG, NGB, TE; Supervision: RNM, H-HK, JG, NGB, TE; Writing – original draft: TE; Writing – review & editing: RNM, LF, JD, DB, SM, AR, H-HK, JG, NGB, TE.

## Data availability

The RNA-Seq data generated during this study have been deposited in the GEO repository (www.ncbi.nlm.nih.gov/geo/) under accession number GSE317084.

## References

1. Ahuja I, Kissen R, Bones AM (2012) Phytoalexins in defense against pathogens. Trends in Plant Science 17: 73–90

2. Bacete L, Mélida H, Miedes E, Molina A (2018) Plant cell wall-mediated immunity: Cell wall changes trigger disease resistance responses. The Plant Journal 93: 614–636

3. Bacete L, Schulz J, Engelsdorf T, Bartosova Z, Vaahtera L, Yan G, Gerhold JM, Tichá T, Øvstebø C, Gigli-Bisceglia N, Johannessen-Starheim S, Margueritat J, Kollist H, Dehoux T, McAdam SAM, Hamann T (2022) THESEUS1 modulates cell wall stiffness and abscisic acid production in *Arabidopsis thaliana*. Proceedings of the National Academy of Sciences 119: e2119258119

4. Bari R, Jones JDG (2009) Role of plant hormones in plant defence responses. Plant Molecular Biology 69: 473–488

5. Bartels S, Boller T (2015) Quo vadis, Pep? Plant elicitor peptides at the crossroads of immunity, stress, and development. Journal of Experimental Botany 66: 5183–5193

6. Bartels S, Lori M, Mbengue M, van Verk M, Klauser D, Hander T, Böni R, Robatzek S, Boller T (2013) The family of Peps and their precursors in Arabidopsis: differential expression and localization but similar induction of pattern-triggered immune responses. Journal of Experimental Botany 64: 5309–5321

7. Bhandari DD, Kim S-J, Brandizzi F (2025) Fortifying the frontier: cell wall modifications during plant immunity. Current Opinion in Plant Biology 88: 102816

8. Birkenbihl RP, Diezel C, Somssich IE (2012) Arabidopsis WRKY33 Is a Key Transcriptional Regulator of Hormonal and Metabolic Responses toward Botrytis cinerea Infection Plant Physiology 159: 266–285

9. Birkenbihl RP, Kracher B, Roccaro M, Somssich IE (2017) Induced Genome-Wide Binding of Three Arabidopsis WRKY Transcription Factors during Early MAMP-Triggered Immunity. The Plant Cell 29: 20–38

10. Bizan J, Sarkar S, Basak AK, Bera S, Goto-Yamada S, Endo K, Tarnawska-Glatt K, Batth R, Bhardwaj K, Mirzaei M, Czerniawski P, Bednarek P, Yamada K (2025) Arabidopsis MYB47 and MYB95 transcription factors regulate jasmonate-inducible ER-body formation. Communications Biology 8: 1377

11. Bücherl CA, Jarsch IK, Schudoma C, Segonzac C, Mbengue M, Robatzek S, MacLean D, Ott T, Zipfel C (2017) Plant immune and growth receptors share common signalling components but localise to distinct plasma membrane nanodomains. eLife 6: e25114

12. Cao J, Yang Q, Zhao Y, Tan S, Li S, Cheng D, Zhang R, Zhang M, Li Z (2025) MYB47 delays leaf senescence by modulating jasmonate pathway via direct regulation of CYP94B3/CYP94C1 expression in Arabidopsis. New Phytologist 246: 2192–2206

13. Chaudhary A, Chen X, Gao J, Leśniewska B, Hammerl R, Dawid C, Schneitz K (2020) The Arabidopsis receptor kinase STRUBBELIG regulates the response to cellulose deficiency. PLOS Genetics 16: e1008433

14. Debnath J, Morton RN, Engelsdorf T, Gigli-Bisceglia N (2026) Plant Cell Wall Remodeling and Peptide Signaling Under Abiotic and Biotic Stress. Plant Communications

15. Denness L, McKenna JF, Segonzac C, Wormit A, Madhou P, Bennett M, Mansfield J, Zipfel C, Hamann T (2011) Cell wall damage-induced lignin biosynthesis is regulated by a reactive oxygen species- and jasmonic acid-dependent process in Arabidopsis. Plant Physiology 156: 1364–1374

16. Engelsdorf T, Gigli-Bisceglia N, Veerabagu M, McKenna JF, Vaahtera L, Augstein F, Van der Does D, Zipfel C, Hamann T (2018) The plant cell wall integrity maintenance and immune signaling systems cooperate to control stress responses in *Arabidopsis thaliana*. Science Signaling 11

17. Engelsdorf T, Horst RJ, Pröls R, Pröschel M, Dietz F, Hückelhoven R, Voll LM (2013) Reduced carbohydrate availability enhances the susceptibility of Arabidopsis toward *Colletotrichum higginsianum*. Plant Physiology 162: 225–238

18. Engelsdorf T, Will C, Hofmann J, Schmitt C, Merritt BB, Rieger L, Frenger MS, Marschall A, Franke RB, Pattathil S, Voll LM (2017) Cell wall composition and penetration resistance against the fungal pathogen *Colletotrichum higginsianum* are affected by impaired starch turnover in Arabidopsis mutants. Journal of Experimental Botany 68: 701–713

19. Feng W, Kita D, Peaucelle A, Cartwright HN, Doan V, Duan Q, Liu M-C, Maman J, Steinhorst L, Schmitz-Thom I, Yvon R, Kudla J, Wu H-M, Cheung AY, Dinneny JR (2018) The FERONIA receptor kinase maintains cell-wall integrity during salt stress through Ca^2+^ signaling. Current Biology 28: 666–675.e665

20. Franck CM, Westermann J, Boisson-Dernier A (2018) Plant malectin-like receptor kinases: From cell wall integrity to immunity and beyond. Annual Review of Plant Biology 69: 301–328

21. Frerigmann H, Glawischnig E, Gigolashvili T (2015) The role of MYB34, MYB51 and MYB122 in the regulation of camalexin biosynthesis in Arabidopsis thaliana. Frontiers in Plant Science 6: 654

22. Gigli Bisceglia N, Savatin DV, Cervone F, Engelsdorf T, De Lorenzo G (2018) Loss of the Arabidopsis Protein Kinases ANPs Affects Root Cell Wall Composition, and Triggers the Cell Wall Damage Syndrome. Frontiers in Plant Science 8

23. Gigli-Bisceglia N, van Zelm E, Huo W, Lamers J, Testerink C (2022) Arabidopsis root responses to salinity depend on pectin modification and cell wall sensing. Development 149: dev200363

24. Glawischnig E (2007) Camalexin. Phytochemistry 68: 401–406

25. Gonneau M, Desprez T, Martin M, Doblas VG, Bacete L, Miart F, Sormani R, Hématy K, Renou J, Landrein B, Murphy E, Van De Cotte B, Vernhettes S, De Smet I, Höfte H (2018) Receptor kinase THESEUS1 is a rapid alkalinization factor 34 receptor in Arabidopsis. Current Biology 28: 2452–2458.e2454

26. Gronnier J, Franck CM, Stegmann M, DeFalco TA, Abarca A, von Arx M, Dünser K, Lin W, Yang Z, Kleine-Vehn J, Ringli C, Zipfel C (2022) Regulation of immune receptor kinase plasma membrane nanoscale organization by a plant peptide hormone and its receptors. eLife 11: e74162

27. Guerreiro J, Marhavý P (2023) Unveiling the intricate mechanisms of plant defense. Frontiers in Plant Physiology 1:1285373

28. Gutierrez R, Lindeboom JJ, Paredez AR, Emons AMC, Ehrhardt DW (2009) Arabidopsis cortical microtubules position cellulose synthase delivery to the plasma membrane and interact with cellulose synthase trafficking compartments. Nature Cell Biology 11: 797

29. Hamann T, Bennett M, Mansfield J, Somerville C (2009) Identification of cell-wall stress as a hexose-dependent and osmosensitive regulator of plant responses. The Plant Journal 57: 1015–1026

30. Hander T, Fernández-Fernández ÁD, Kumpf RP, Willems P, Schatowitz H, Rombaut D, Staes A, Nolf J, Pottie R, Yao P, Gonçalves A, Pavie B, Boller T, Gevaert K, Van Breusegem F, Bartels S, Stael S (2019) Damage on plants activates Ca2+-dependent metacaspases for release of immunomodulatory peptides. Science 363: eaar7486

31. Hdedeh O, Mercier C, Poitout A, Martinière A, Zelazny E (2025) Membrane nanodomains to shape plant cellular functions and signaling. New Phytologist 245: 1369–1385

32. He Y, Xu J, Wang X, He X, Wang Y, Zhou J, Zhang S, Meng X (2019) The Arabidopsis Pleiotropic Drug Resistance Transporters PEN3 and PDR12 Mediate Camalexin Secretion for Resistance to Botrytis cinerea. The Plant Cell 31: 2206–2222

33. Hématy K, Sado P-E, Van Tuinen A, Rochange S, Desnos T, Balzergue S, Pelletier S, Renou J-P, Höfte H (2007) A receptor-like kinase mediates the response of Arabidopsis cells to the inhibition of cellulose synthesis. Current Biology 17: 922–931

34. Hou S, Liu D, Huang S, Luo D, Liu Z, Xiang Q, Wang P, Mu R, Han Z, Chen S, Chai J, Shan L, He P (2021) The Arabidopsis MIK2 receptor elicits immunity by sensing a conserved signature from phytocytokines and microbes. Nature Communications 12: 5494

35. Jaillais Y, Bayer E, Bergmann DC, Botella MA, Boutté Y, Bozkurt TO, Caillaud M-C, Germain V, Grossmann G, Heilmann I, Hemsley PA, Kirchhelle C, Martinière A, Miao Y, Mongrand S, Müller S, Noack LC, Oda Y, Ott T, Pan X, Pleskot R, Potocky M, Robert S, Rodriguez CS, Simon-Plas F, Russinova E, Van Damme D, Van Norman JM, Weijers D, Yalovsky S, Yang Z, Zelazny E, Gronnier J (2024) Guidelines for naming and studying plasma membrane domains in plants. Nature Plants 10: 1172–1183

36. Jolivet M-D, Deroubaix AF, Boudsocq M, Abel NB, Rocher M, Robbe T, Wattelet-Boyer V, Huard J, Lefebvre D, Lu Y-J, Day B, Saias G, Ahmed J, Cotelle V, Giovinazzo N, Gallois J-L, Yamaji Y, German-Retana S, Gronnier J, Ott T, Mongrand S, Germain V (2025) Interdependence of plasma membrane nanoscale dynamics of a kinase and its cognate substrate underlies Arabidopsis response to viral infection. eLife 12: RP90309

37. Knight H, Trewavas AJ, Knight MR (1996) Cold calcium signaling in Arabidopsis involves two cellular pools and a change in calcium signature after acclimation. The Plant Cell 8: 489–503

38. Knight MR, Campbell AK, Smith SM, Trewavas AJ (1991) Transgenic plant aequorin reports the effects of touch and cold-shock and elicitors on cytoplasmic calcium. Nature 352: 524–526

39. Konopka CA, Bednarek SY (2008) Variable-angle epifluorescence microscopy: a new way to look at protein dynamics in the plant cell cortex. The Plant Journal 53: 186–196

40. Laine RF, Heil HS, Coelho S, Nixon-Abell J, Jimenez A, Wiesner T, Martínez D, Galgani T, Régnier L, Stubb A, Follain G, Webster S, Goyette J, Dauphin A, Salles A, Culley S, Jacquemet G, Hajj B, Leterrier C, Henriques R (2023) High-fidelity 3D live-cell nanoscopy through data-driven enhanced super-resolution radial fluctuation. Nature Methods 20: 1949–1956

41. Li Y, Liu K, Tong G, Xi C, Liu J, Zhao H, Wang Y, Ren D, Han S (2021) MPK3/MPK6-mediated phosphorylation of ERF72 positively regulates resistance to Botrytis cinerea through directly and indirectly activating the transcription of camalexin biosynthesis enzymes. Journal of Experimental Botany 73: 413–428

42. Liu M-CJ, Yeh F-LJ, Yvon R, Simpson K, Jordan S, Chambers J, Wu H-M, Cheung AY (2024) Extracellular pectin-RALF phase separation mediates FERONIA global signaling function. Cell 187: 312–330.e322

43. Mao G, Meng X, Liu Y, Zheng Z, Chen Z, Zhang S (2011) Phosphorylation of a WRKY Transcription Factor by Two Pathogen-Responsive MAPKs Drives Phytoalexin Biosynthesis in Arabidopsis. The Plant Cell 23: 1639–1653

44. Marquis V, Smirnova E, Graindorge S, Delcros P, Villette C, Zumsteg J, Heintz D, Heitz T (2022) Broad-spectrum stress tolerance conferred by suppressing jasmonate signaling attenuation in Arabidopsis JASMONIC ACID OXIDASE mutants. The Plant Journal 109: 856–872

45. Martinière A, Lavagi I, Nageswaran G, Rolfe DJ, Maneta-Peyret L, Luu D-T, Botchway SW, Webb SED, Mongrand S, Maurel C, Martin-Fernandez ML, Kleine-Vehn J, Friml J, Moreau P, Runions J (2012) Cell wall constrains lateral diffusion of plant plasma-membrane proteins. Proceedings of the National Academy of Sciences 109: 12805–12810

46. McKenna JF, Rolfe DJ, Webb SED, Tolmie AF, Botchway SW, Martin-Fernandez ML, Hawes C, Runions J (2019) The cell wall regulates dynamics and size of plasma-membrane nanodomains in Arabidopsis. Proceedings of the National Academy of Sciences 116: 12857–12862

47. Merz D, Richter J, Gonneau M, Sanchez-Rodriguez C, Eder T, Sormani R, Martin M, Hématy K, Höfte H, Hauser M-T (2017) T-DNA alleles of the receptor kinase THESEUS1 with opposing effects on cell wall integrity signaling. Journal of Experimental Botany 68: 4583–4593

48. Mielke S, Zimmer M, Meena MK, Dreos R, Stellmach H, Hause B, Voiniciuc C, Gasperini D (2021) Jasmonate biosynthesis arising from altered cell walls is prompted by turgor-driven mechanical compression. Science Advances 7: eabf0356

49. Millard PS, Weber K, Kragelund BB, Burow M (2019) Specificity of MYB interactions relies on motifs in ordered and disordered contexts. Nucleic Acids Research 47: 9592–9608

50. Molina A, Miedes E, Bacete L, Rodríguez T, Mélida H, Denancé N, Sánchez-Vallet A, Rivière M-P, López G, Freydier A, Barlet X, Pattathil S, Hahn M, Goffner D (2021) Arabidopsis cell wall composition determines disease resistance specificity and fitness. Proceedings of the National Academy of Sciences 118: e2010243118

51. Monsalvo I, Parasecolo L, Pullano S, Lin J, Shahabi A, Ly M, Kwon H, Mathur K, Rodrillo KAM, Ifa DR, Kovinich N (2025) ANAC042 Regulates the Biosynthesis of Conserved- and Lineage-Specific Phytoalexins in Arabidopsis. International Journal of Molecular Sciences 26: 3683

52. Mucha S, Heinzlmeir S, Kriechbaumer V, Strickland B, Kirchhelle C, Choudhary M, Kowalski N, Eichmann R, Hückelhoven R, Grill E, Kuster B, Glawischnig E (2019) The Formation of a Camalexin Biosynthetic Metabolon. The Plant Cell 31: 2697–2710

53. Müller TM, Böttcher C, Morbitzer R, Götz CC, Lehmann J, Lahaye T, Glawischnig E (2015) TRANSCRIPTION ACTIVATOR-LIKE EFFECTOR NUCLEASE-Mediated Generation and Metabolic Analysis of Camalexin-Deficient *cyp71a12 cyp71a13* Double Knockout Lines. Plant Physiology 168: 849–858

54. Munzert KS, Engelsdorf T (2025) Plant cell wall structure and dynamics in plant–pathogen interactions and pathogen defence. Journal of Experimental Botany 76: 228–242

55. Nakagawa Y, Katagiri T, Shinozaki K, Qi Z, Tatsumi H, Furuichi T, Kishigami A, Sokabe M, Kojima I, Sato S, Kato T, Tabata S, Iida K, Terashima A, Nakano M, Ikeda M, Yamanaka T, Iida H (2007) Arabidopsis plasma membrane protein crucial for Ca2+ influx and touch sensing in roots. Proceedings of the National Academy of Sciences 104: 3639–3644

56. Nakaminami K, Okamoto M, Higuchi-Takeuchi M, Yoshizumi T, Yamaguchi Y, Fukao Y, Shimizu M, Ohashi C, Tanaka M, Matsui M, Shinozaki K, Seki M, Hanada K (2018) AtPep3 is a hormone-like peptide that plays a role in the salinity stress tolerance of plants. Proceedings of the National Academy of Sciences 115: 5810–5815

57. Narusaka Y, Narusaka M, Park P, Kubo Y, Hirayama T, Seki M, Shiraishi T, Ishida J, Nakashima M, Enju A, Sakurai T, Satou M, Kobayashi M, Shinozaki K (2004) RCH1, a locus in Arabidopsis that confers resistance to the hemibiotrophic fungal pathogen *Colletotrichum higginsianum*. Molecular Plant-Microbe Interactions 17: 749–762

58. Nawrath C, Métraux J-P (1999) Salicylic Acid Induction–Deficient Mutants of Arabidopsis Express PR-2 and PR-5 and Accumulate High Levels of Camalexin after Pathogen Inoculation. The Plant Cell 11: 1393–1404

59. Nomura H, Komori T, Kobori M, Nakahira Y, Shiina T (2008) Evidence for chloroplast control of external Ca2+-induced cytosolic Ca2+ transients and stomatal closure. The Plant Journal 53: 988–998

60. Nomura H, Komori T, Uemura S, Kanda Y, Shimotani K, Nakai K, Furuichi T, Takebayashi K, Sugimoto T, Sano S, Suwastika IN, Fukusaki E, Yoshioka H, Nakahira Y, Shiina T (2012) Chloroplast-mediated activation of plant immune signalling in Arabidopsis. Nature Communications 3: 926

61. Novaković L, Guo T, Bacic A, Sampathkumar A, Johnson KL (2018) Hitting the Wall—Sensing and Signaling Pathways Involved in Plant Cell Wall Remodeling in Response to Abiotic Stress. Plants 7: 89

62. O’Connell R, Herbert C, Sreenivasaprasad S, Khatib M, Esquerré-Tugayé M-T, Dumas B (2004) A novel Arabidopsis-*Colletotrichum* pathosystem for the molecular dissection of plant-fungal interactions. Molecular Plant-Microbe Interactions 17: 272–282

63. Ortiz-Morea FA, Liu J, Shan L, He P (2022) Malectin-like receptor kinases as protector deities in plant immunity. Nature Plants 8: 27–37

64. Paredez AR, Somerville CR, Ehrhardt DW (2006) Visualization of cellulose synthase demonstrates functional association with microtubules. Science 312: 1491–1495

65. Pastorczyk M, Kosaka A, Piślewska-Bednarek M, López G, Frerigmann H, Kułak K, Glawischnig E, Molina A, Takano Y, Bednarek P (2020) The role of CYP71A12 monooxygenase in pathogen-triggered tryptophan metabolism and Arabidopsis immunity. New Phytologist 225: 400–412

66. Perraki A, Gronnier J, Gouguet P, Boudsocq M, Deroubaix A-F, Simon V, German-Retana S, Legrand A, Habenstein B, Zipfel C, Bayer E, Mongrand S, Germain V (2018) REM1.3’s phospho-status defines its plasma membrane nanodomain organization and activity in restricting PVX cell-to-cell movement. PLOS Pathogens 14: e1007378

67. Reckleben L, Morton RN, Munzert-Eberlein KS, Thiel S, Rautengarten C, Tan V, Harres KL, Zanoni L, Ebert B, Engelsdorf T (2025) Galactan mobilization during carbon starvation compromises plant cell wall-mediated resistance to fungal infection. The Plant Journal 123: e70438

68. Rhodes J, Yang H, Moussu S, Boutrot F, Santiago J, Zipfel C (2021) Perception of a divergent family of phytocytokines by the Arabidopsis receptor kinase MIK2. Nature Communications 12: 705

69. Rohr L, Ehinger A, Rausch L, Glöckner Burmeister N, Meixner AJ, Gronnier J, Harter K, Kemmerling B, zur Oven-Krockhaus S (2024) OneFlowTraX: a user-friendly software for super-resolution analysis of single-molecule dynamics and nanoscale organization. Frontiers in Plant Science 15: 1358935

70. Saga H, Ogawa T, Kai K, Suzuki H, Ogata Y, Sakurai N, Shibata D, Ohta D (2012) Identification and characterization of ANAC042, a transcription factor family gene involved in the regulation of camalexin biosynthesis in Arabidopsis. Molecular Plant-Microbe Interactions 25: 684–696

71. Schindelin J, Arganda-Carreras I, Frise E, Kaynig V, Longair M, Pietzsch T, Preibisch S, Rueden C, Saalfeld S, Schmid B, Tinevez J-Y, White DJ, Hartenstein V, Eliceiri K, Tomancak P, Cardona A (2012) Fiji: an open-source platform for biological-image analysis. Nature Methods 9: 676–682

72. Schmidt A, Mächtel R, Ammon A, Engelsdorf T, Schmitz J, Maurino VG, Voll LM (2020) Reactive oxygen species dosage in Arabidopsis chloroplasts can improve resistance towards Colletotrichum higginsianum by the induction of WRKY33. New Phytologist 226: 189–204

73. Schoenaers S, Vissenberg K (2025) Overlooked aspects of CrRLK1L–RALF signaling. New Phytologist

74. Shen W, Liu J, Li J-F (2019) Type-II metacaspases mediate the processing of plant elicitor peptides in Arabidopsis. Molecular Plant 12: 1524–1533

75. Tanaka K, Choi J, Stacey G (2013) Aequorin Luminescence-Based Functional Calcium Assay for Heterotrimeric G-Proteins in Arabidopsis. *In* MP Running, ed, G Protein-Coupled Receptor Signaling in Plants: Methods and Protocols. Humana Press, Totowa, NJ, pp 45–54

76. Tang J, Han Z, Sun Y, Zhang H, Gong X, Chai J (2015) Structural basis for recognition of an endogenous peptide by the plant receptor kinase PEPR1. Cell Research 25: 110–120

77. Vaahtera L, Schulz J, Hamann T (2019) Cell wall integrity maintenance during plant development and interaction with the environment. Nature Plants 5: 924–932

78. Van der Does D, Boutrot F, Engelsdorf T, Rhodes J, McKenna JF, Vernhettes S, Koevoets I, Tintor N, Veerabagu M, Miedes E, Segonzac C, Roux M, Breda AS, Hardtke CS, Molina A, Rep M, Testerink C, Mouille G, Höfte H, Hamann T, Zipfel C (2017) The Arabidopsis leucine-rich repeat receptor kinase MIK2/LRR-KISS connects cell wall integrity sensing, root growth and response to abiotic and biotic stresses. PLOS Genetics 13: e1006832

79. Völkner C, Holzner LJ, Day PM, Ashok AD, Vries Jd, Bölter B, Kunz H-H (2021) Two plastid POLLUX ion channel-like proteins are required for stress-triggered stromal Ca2+release. Plant Physiology 187: 2110–2125

80. Voll LM, Zell MB, Engelsdorf T, Saur A, Wheeler MG, Drincovich MF, Weber APM, Maurino VG (2012) Loss of cytosolic NADP-malic enzyme 2 in Arabidopsis thaliana is associated with enhanced susceptibility to *Colletotrichum higginsianum*. New Phytologist 195: 189–202

81. von Arx M, Xhelilaj K, Schulz P, zur Oven-Krockhaus S, Gronnier J (2024) Photochromic reversion enables long-term tracking of single molecules in living plants. bioRxiv: 2024.2004.2010.585335

82. Weinl S, Held K, Schlücking K, Steinhorst L, Kuhlgert S, Hippler M, Kudla J (2008) A plastid protein crucial for Ca2+-regulated stomatal responses. New Phytologist 179: 675–686

83. Wolf S (2022) Cell Wall Signaling in Plant Development and Defense. Annual Review of Plant Biology 73: 323–353

84. Yamaguchi Y, Huffaker A, Bryan AC, Tax FE, Ryan CA (2010) PEPR2 Is a Second Receptor for the Pep1 and Pep2 Peptides and Contributes to Defense Responses in Arabidopsis The Plant Cell 22: 508–522

85. Yamaguchi Y, Pearce G, Ryan CA (2006) The cell surface leucine-rich repeat receptor for *At*Pep1, an endogenous peptide elicitor in Arabidopsis, is functional in transgenic tobacco cells. Proceedings of the National Academy of Sciences 103: 10104–10109

86. Zhai K, Rhodes J, Zipfel C (2024) A peptide-receptor module links cell wall integrity sensing to pattern-triggered immunity. Nature Plants 10: 2027–2037

87. Zhang C, Wu Y, Liu J, Song B, Yu Z, Li J-F, Yang C, Lai J (2025) SUMOylation controls peptide processing to generate damage-associated molecular patterns in Arabidopsis. Developmental Cell 60: 696–705.e694

88. Zhang Z, Gigli-Bisceglia N, Li W, Li S, Wang J, Liu J, Testerink C, Guo Y (2024) SCOOP10 and SCOOP12 peptides act through MIK2 receptor-like kinase to antagonistically regulate Arabidopsis leaf senescence. Molecular Plant 17: 1805–1819

89. Zhou J, Mu Q, Wang X, Zhang J, Yu H, Huang T, He Y, Dai S, Meng X (2022) Multilayered synergistic regulation of phytoalexin biosynthesis by ethylene, jasmonate, and MAPK signaling pathways in Arabidopsis. The Plant Cell 34: 3066–3087

90. Zhou J, Wang X, He Y, Sang T, Wang P, Dai S, Zhang S, Meng X (2020) Differential phosphorylation of the transcription factor WRKY33 by the protein kinases CPK5/CPK6 and MPK3/MPK6 cooperatively regulates camalexin biosynthesis in Arabidopsis. The Plant Cell 32: 2621–2638

