## Supplemental Figures for "Cell wall integrity and elicitor peptide signaling modulate jasmonic acid-mediated camalexin production in Arabidopsis"

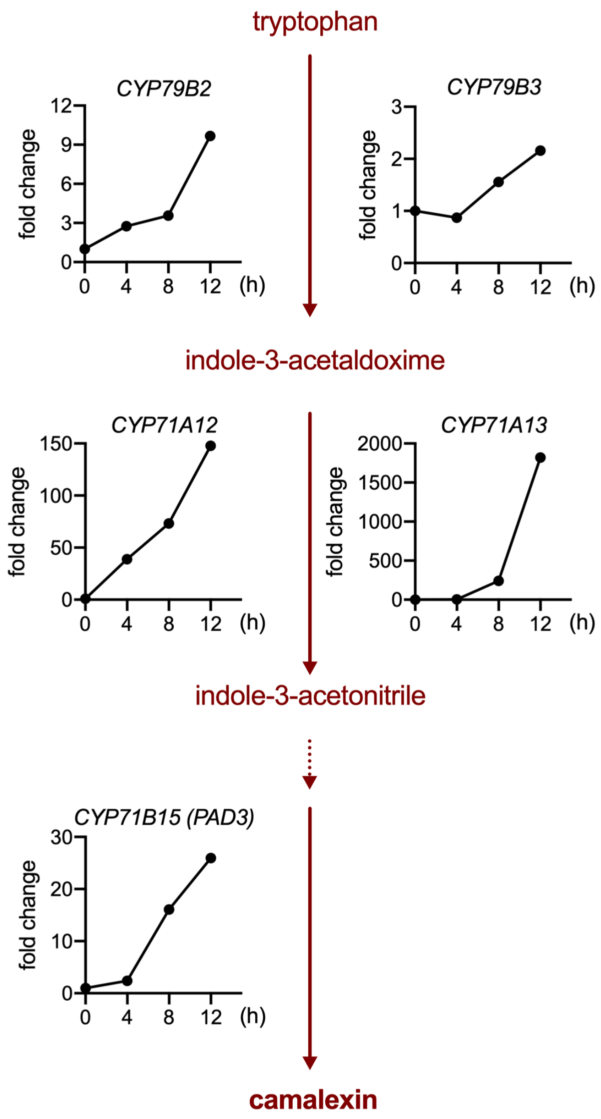


**Figure S1: Published transcriptomics data suggest activation of the camalexin biosynthesis pathway by isoxaben.**
Relative gene expression of *CYP79B2*, *CAP79B3*, *CYP71A12*, *CYP71A13* and *CYP71B15*/*PAD3* after 0, 4, 8 and 12 h of isoxaben (ISX) treatment compared to t = 0 h (fold change). Microarray data were taken from Hamann et al. (2009). Graphs are positioned along the camalexin biosynthesis pathway according to the enzymatic functions encoded by the genes.

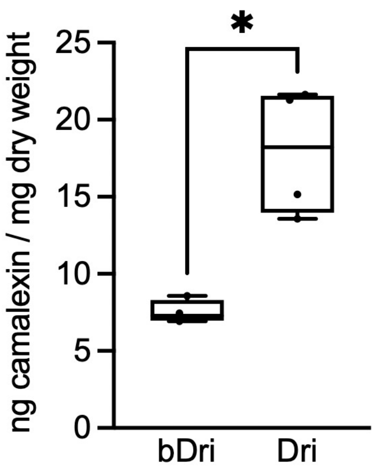


**Figure S2: Camalexin accumulates after Driselase treatment.**
Camalexin amount in Col-0 seedlings was determined at 8 h after treatment with boiled Driselase (bDri) or Driselase (Dri) (n=4). Asterisks indicate statistically significant differences according to a Student's t-test (*$p < 0.05$).

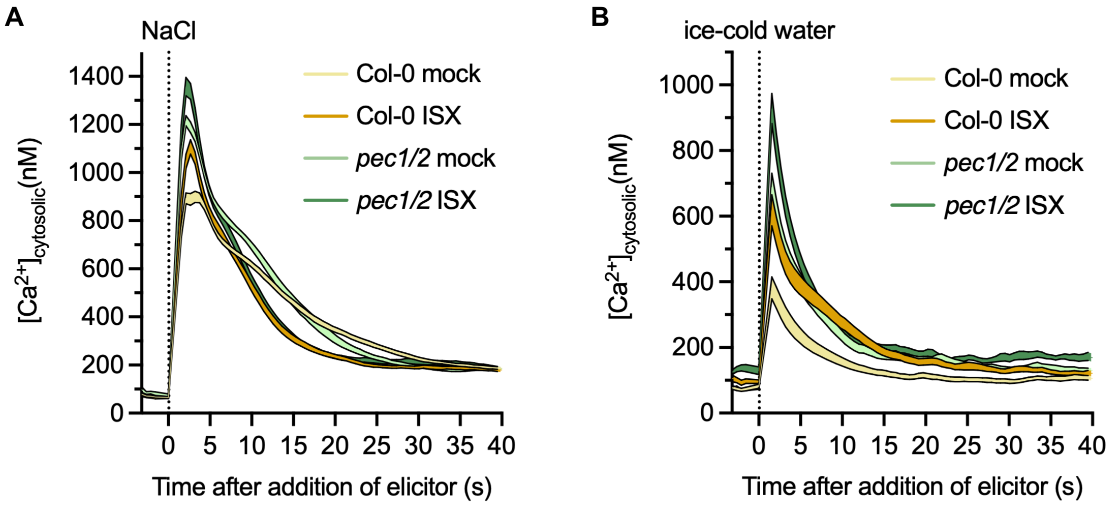


**Figure S3: CWD caused by ISX treatment increases salt- and cold-induced calcium transients**
Cytosolic $Ca^{2+}$ concentration in Col-0 and *pec1/2* seedlings after elicitation with (A) 200 mM NaCl (n=16) and (B) ice-cold water (n=17-24). Prior to elicitation, seedlings were treated for 4.5 h with mock or ISX. The shaded area indicates the SEM.

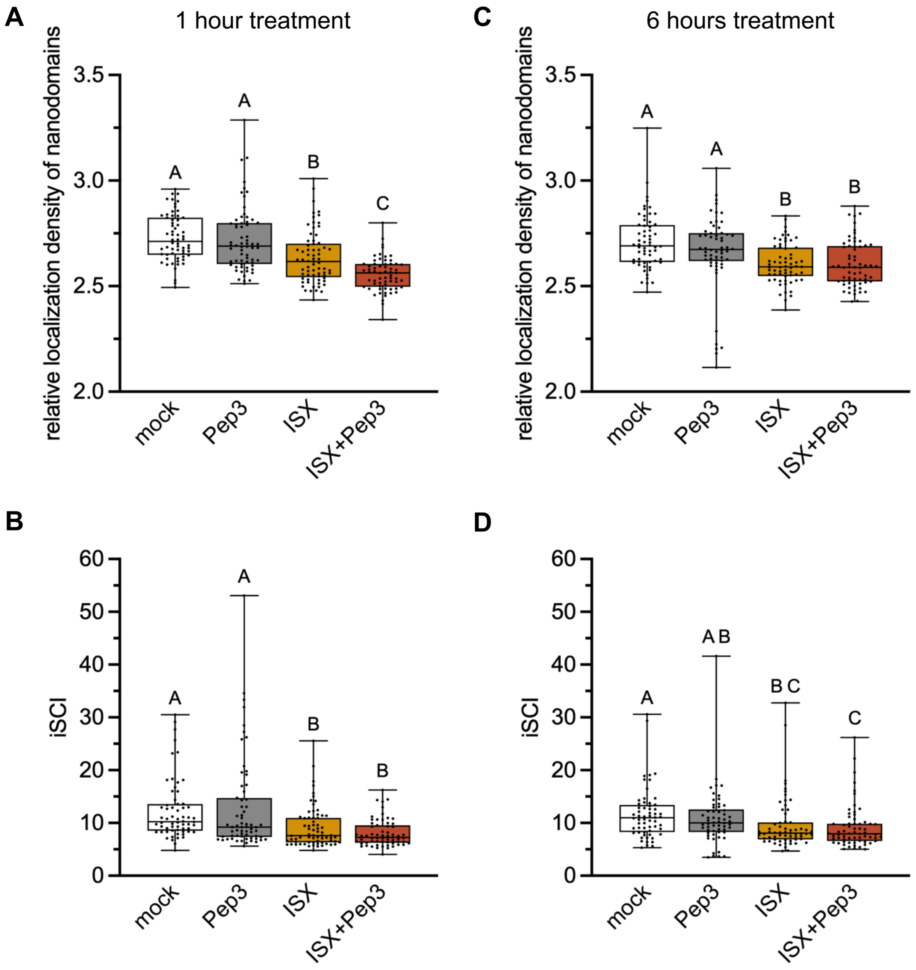


**Figure S4: THE1-GFP plasma membrane nanodomain organization after ISX, Pep3 and combined treatments.**

THE1-GFP nanodomain organization was analyzed with variable angle total internal reflection microscopy (VA-TIRFM) in combination with enhanced-super resolution radial fluctuation analyses (eSRRF) in seedling hypocotyls at (A, B) 1 h and (C, D) 6 h after treatment with mock, Pep3, isoxaben (ISX) and ISX+Pep3. (A, C) relative localization density of nanodomains and (B, D) mean image wide spatial clustering index (iSCI) per cell were recorded in three pooled experiments with 4-8 cells from three seedlings per experiment (n=58-69). Different letters indicate statistically significant differences according to Kruskal-Wallis and Dunn’s test (α = 0.05).

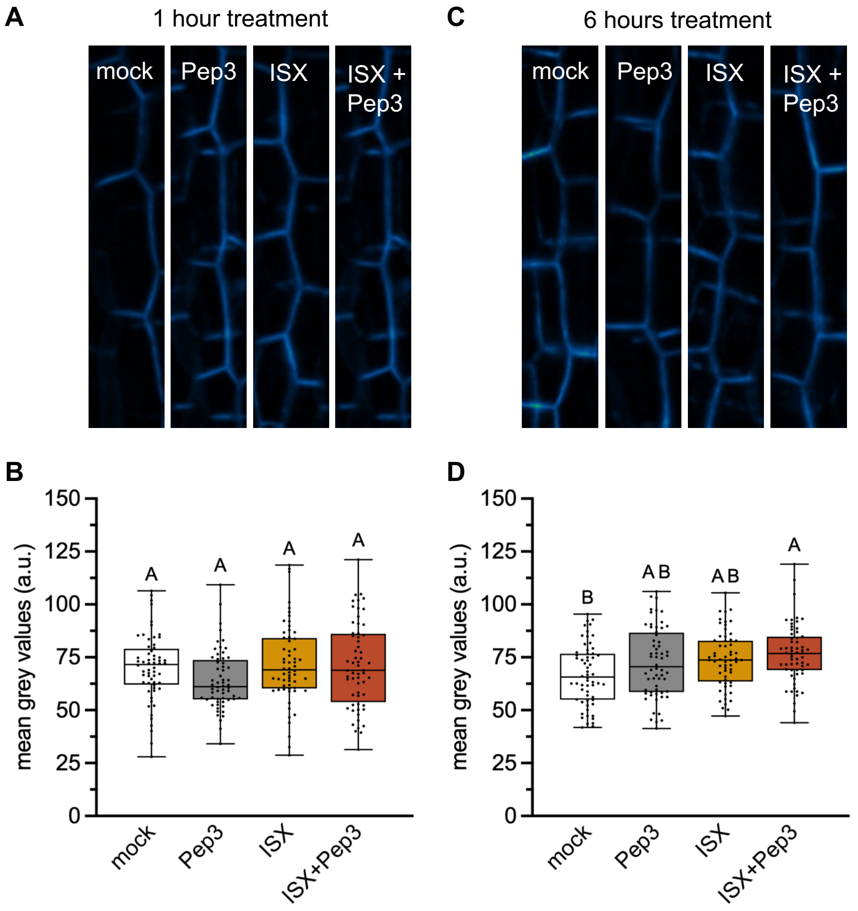


**Figure S5: THE1-GFP localization to the plasma membrane after ISX, Pep3 and combined treatments.**

THE1-GFP fluorescence intensity was determined in seedling hypocotyls at (A, B) 1 h and (C, D) 6 h after treatment with mock, Pep3, isoxaben (ISX) and ISX+Pep3. (A, C) Representative images taken with confocal laser scanning microscopy. (B, D) Mean grey values recorded in two pooled experiments with 10 cells from three seedlings per experiment (n=60). Different letters indicate statistically significant differences according to Kruskal-Wallis and Dunn's test ($\alpha = 0.05$).

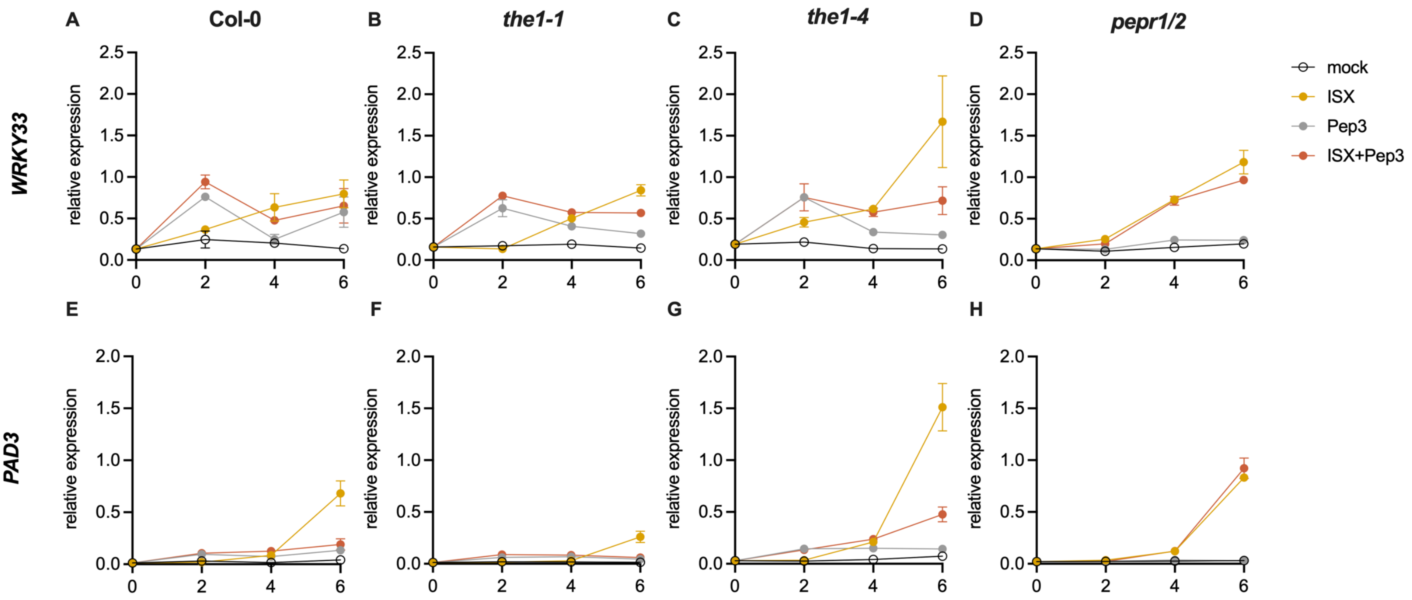


**Figure S6: Time-course analysis of *WRKY33* and *PAD3* expression after Pep3 and ISX treatments.** Relative expression of (A-D) *WRKY33* and (E-H) *PAD3* was determined in (A, E) Col-0, (B, F) *the1-1*, (C, G) *the1-4* and (D, H) *pepr1 pepr2* at 0, 2, 4 and 6 h after treatment with mock, Pep3, isoxaben (ISX) and ISX+Pep3 via quantitative RT-PCR. Values are relative to *ACT2* expression and error bars represent the SEM (n=3). Please note that the values for t = 4 h and 6 h are also shown in Figure 4 A-D.

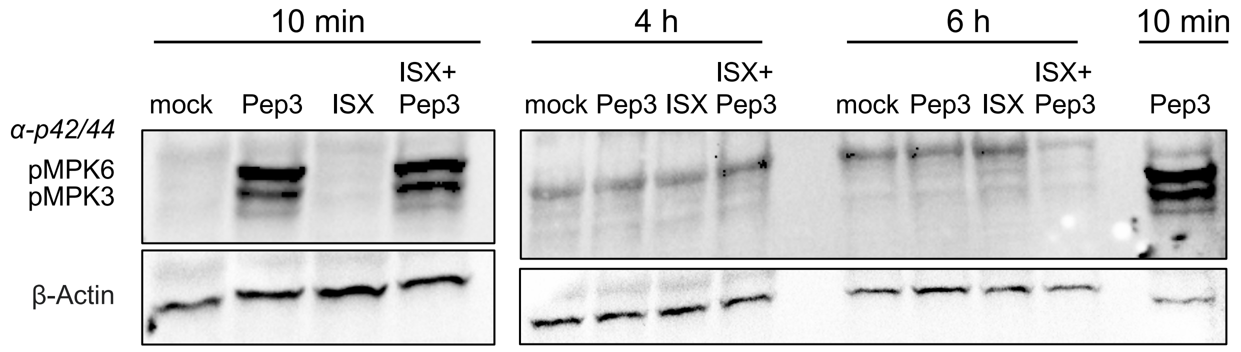


**Figure S7: Investigation of MPK phosphorylation after ISX and Pep3 treatments.**
Phosphorylation of MPK3 and MPK6 was determined in Col-0 seedlings at 10 min, 4 h and 6 h after treatment with mock, Pep3, isoxaben (ISX) and ISX+Pep3 by immunoblotting. Equal loading was checked with an anti-β-Actin antibody. Similar results were obtained in three biological replicates.

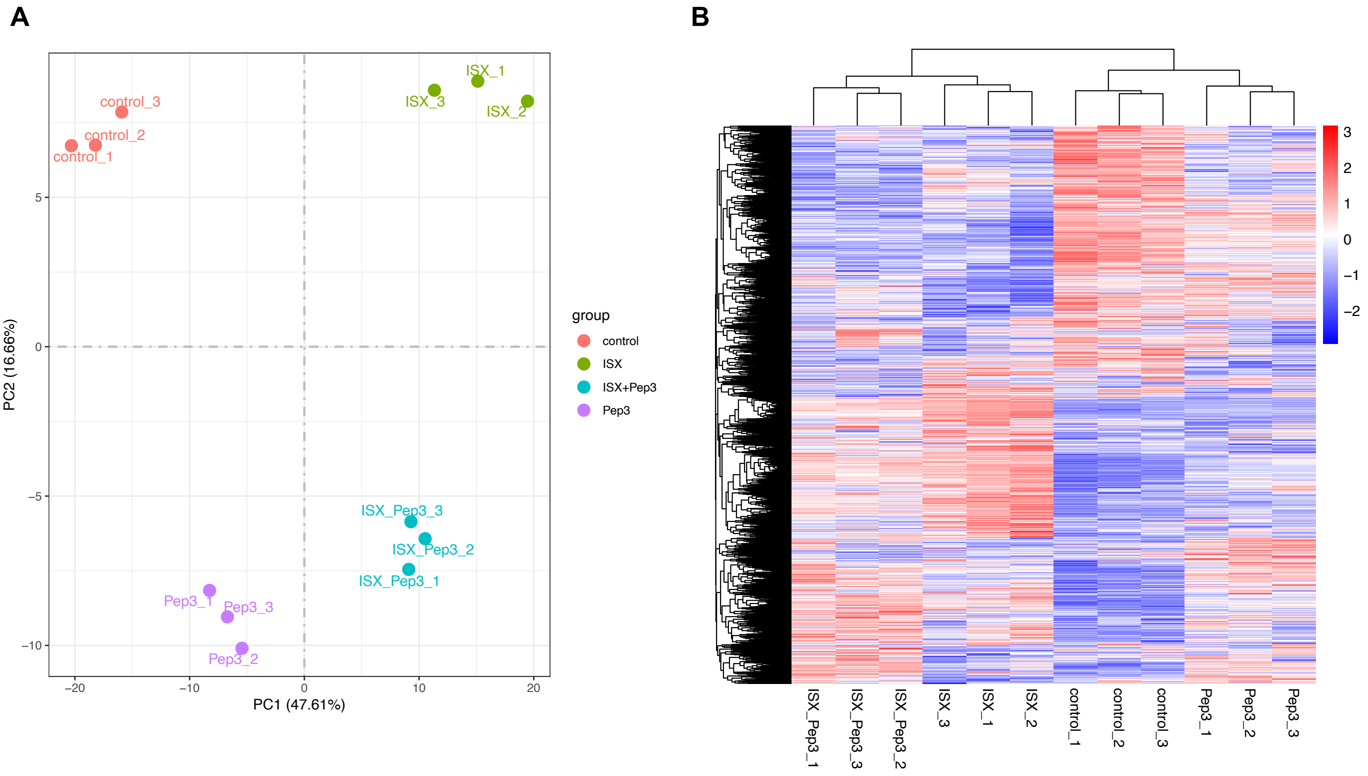


**Figure S8: Principal component and hierarchical cluster analyses of RNA-Sequencing data.** (A) Principal component analysis (PCA) of RNA-Sequencing data for Col-0 seedlings treated with mock, Pep3, isoxaben (ISX) and ISX+Pep3 for 6 h shows grouping of individual replicates (n=3) and main effects of treatments on gene expression. (B) Hierarchical cluster analysis displays similarities in gene expression after the different treatments and identifies sub-clusters. Heat map shows z-score of normalized data.

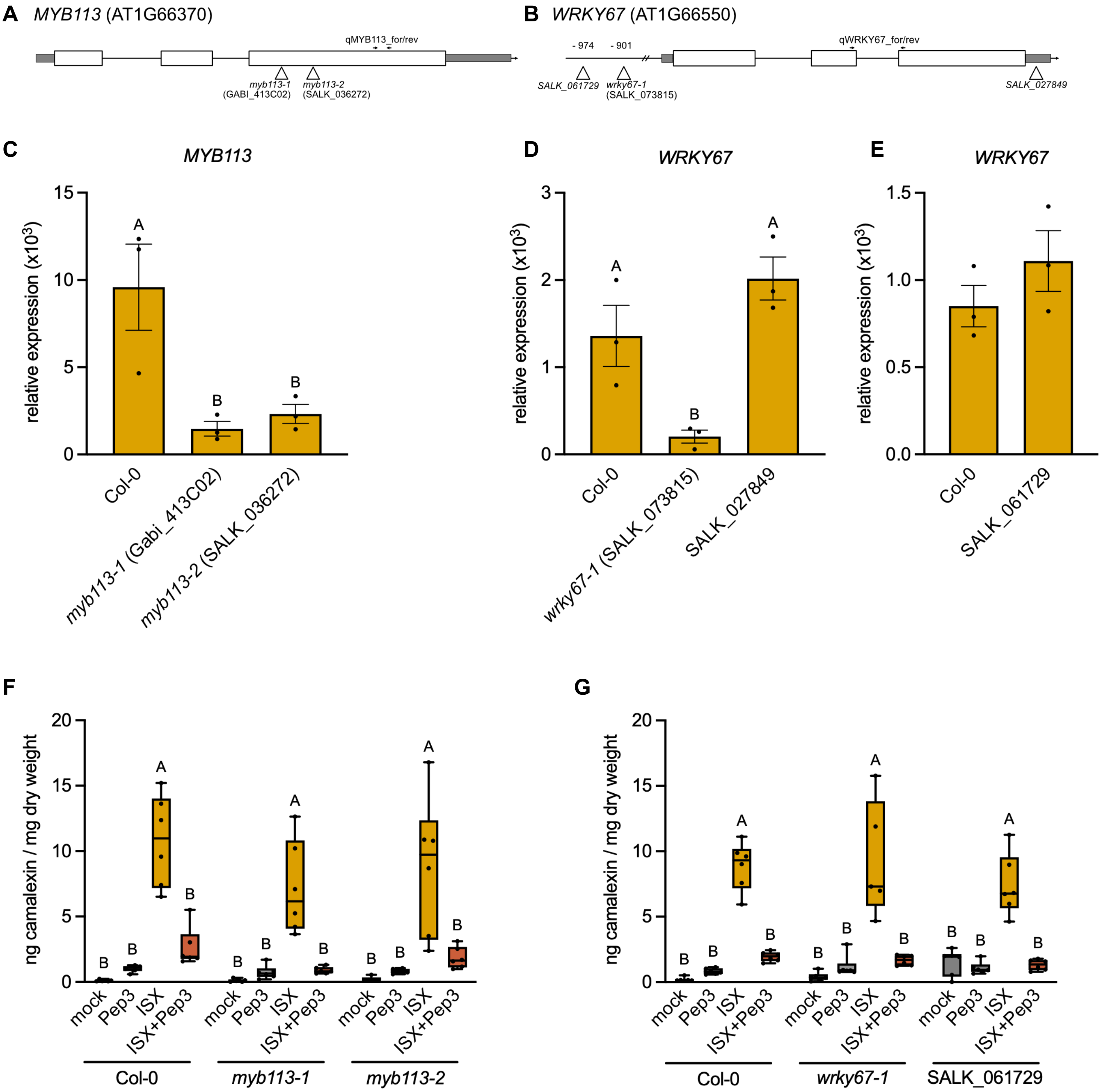


**Figure S9: Genetic characterization and camalexin analysis of *myb113* and *wrky67* mutants.** (A,B) Sketches of the (A) *MYB113* and (B) *WRKY67* genes indicate untranslated regions (grey boxes), exons (white boxes), introns (lines), T-DNA insertion sites (triangles), and primers used for quantitative RT-PCR analysis (arrows). (C) Relative expression of *MYB113* was determined in Col-0, *myb113-1* (Gabi_413C02) and *myb113-2* (SALK_036272) seedlings after 6 h treatment with isoxaben (ISX) via quantitative RT-PCR. (D,E) Relative expression of *WRKY67* was determined in Col-0, *wrky67-1* (SALK_073815), SALK_027849 and SALK_061729 seedlings after 6 h treatment with ISX via quantitative RT-PCR. Values in C-E are relative to *ACT2* expression and error bars represent the SEM (n=3). Different letters indicate statistically significant differences according to one-way ANOVA and Holm-Sidak's multiple comparisons test ($\alpha$ = 0.05). (F,G) Camalexin amount in (F) Col-0, *myb113-1* and *myb113-2,* and in (G) Col-0, *wrky67-1* and SALK_061729 seedlings at 8 h after mock, Pep3, ISX and ISX+Pep3 treatment (n=6). Different letters indicate statistically significant differences according to two-way ANOVA and Holm-Sidak's multiple comparisons test ($\alpha$ = 0.05).
