## Supplemental Tables for "Cell wall integrity and elicitor peptide signaling modulate jasmonic acid-mediated camalexin production in Arabidopsis"

**Table S1.** Arabidopsis genotypes used in this study.

| Genotype | AGI | Stock code | Reference |
|---|---|---|---|
| *the1-1* | At5g54380 | EMS mutant | Hématy et al. (2007) |
| *the1-4* | At5g54380 | SAIL_683_H03 | Guo et al. (2009) |
| *pad3-1* | At3g26830 | EMS mutant | Glazebrook and Ausubel (1994) |
| *the1-1 pad3-1* | At5g54380, At3g26830 | EMS mutant | this study |
| *cyp71a12* | At2g30750 | GABI_127H03 | Millet et al. (2010) |
| *cyp71a13* | At2g30770 | SALK_105136 | Nafisi et al. (2007) |
| *cyp71a12 cyp71a13* | At2g30750, At2g30770 | TALEN SALK_105136 | Müller et al. (2015) |
| *cas-1* | At5g23060 | SALK_070416 | Weinl et al. (2008) |
| *mca1-null* | At4g35920 | n/a | Nakagawa et al. (2007) |
| *pepr1-1 pepr2-1* | At1g73080, At1g17750 | SALK_059281, SALK_036564 | Yamaguchi et al. (2010) |
| *pec1-1 pec2-1* | At5g02940, At5g43745 | SAIL_300_A10, SALK_102200 | Völkner et al. (2021) |
| *wrky33-2* | At5g54380, At2g38470 | GABI_324B11 | Mao et al. (2011) |
| *the1-1 wrky33-2* | At5g54380, At2g38470 | EMS mutant, GABI_324B11 | this study |
| *the1-4 wrky33-2* | At5g54380, At2g38470 | SAIL_683_H03, GABI_324B11 | this study |
| *mpk3-1* | At3g45640 | SALK_151594 | Wang et al. (2007) |
| *mpk6-3* | At2g43790 | SALK_127507 | Liu and Zhang (2004) |
| *pTHE1::THE1-GFP (the1-1)* | At5g54380 | n/a | Chaudhary et al. (2021) |
| *myb47-2* | At1g18710 | SALK_200360 | Renard et al. (2020) |
| *myb47-1 myb95* | At1g18710, At1g74430 | SM_3_38234, GABI_314F05 | Zhang et al. (2019) |
| *myb113-1* | At1g66370 | GABI_413C02 | this study |
| *myb113-2* | At1g66370 | SALK_036272 | this study |
| *wrky67-1* | At1g66550 | SALK_073815 | this study |
| *SALK_061729* | At1g66550 | SALKseq_061729 | this study |
| *SALK_027849* | At1g66550 | SALK_027849C | this study |
| *aos* | At5g42650 | n/a | Park et al. (2002) |

**Table S2.** Primers used in this study.

| Name | Method | Sequence (5'-3') | Reference |
|---|---|---|---|
| qACT2_for | qRT-PCR | CTTGCACCAAGCAGCATGAA | Czechowski et al. (2005) |
| qACT2_rev | | CCGATCCAGACACTGTACTTCCTT | |
| qWRKY33_for | qRT-PCR | TAGAGCAAAGGAAGAACCAAACG | Schmidt et al. (2020) |
| qWRKY33_rev | | TGGTGGCAGAAATGTACAAAAGG | |
| qPAD3_for | qRT-PCR | TCGCTGGCATAACACTATGG | Giovannoni et al. (2021) |
| qPAD3_rev | | TTGGGAGCAAGAGTGGAGT | |
| qCYP71A13_for | qRT-PCR | ATTCGGATCAGGGAGAAGGATA | Birkenbihl et al. (2012) |
| qCYP71A13_rev | | CGATACCAATGGCTTCAGTTAGAT | |
| qPROPEP3_for | qRT-PCR | CAACGATGGAGAATCTCAGA | Engelsdorf et al. (2018) |
| qPROPEP3_rev | | CTAATTGTGTTTGCCTCCTTT | |
| qMYB47_for | qRT-PCR | GCTTACATCAACGAGCATGG | this study |
| qMYB47_rev | | TCCACATCTCTGCAAACCAG | |
| qMYB95_for | qRT-PCR | TCTCCTACACCGAGCCCTAC | this study |
| qMYB95_rev | | GCGAGCTTGTTAAGGAAACG | |
| qMYB113_for | qRT-PCR | TACTGGAGGCAGATGTGTTGG | this study |
| qMYB113_rev | | AAGTCAAGCGGTAAGGTCACA | |
| qWRKY67_for | qRT-PCR | AAAGCCTCAGCGCACAAAAG | this study |
| qWRKY67_rev | | TTGATCTTCTGCACCCGCTT | |

### Supplemental References


**Birkenbihl RP, Diezel C, Somssich IE** (2012) Arabidopsis WRKY33 Is a Key Transcriptional Regulator of Hormonal and Metabolic Responses toward Botrytis cinerea Infection Plant Physiology **159:** 266-285

**Chaudhary A, Chen X, Leśniewska B, Boikine R, Gao J, Wolf S, Schneitz K** (2021) Cell wall damage attenuates root hair patterning and tissue morphogenesis mediated by the receptor kinase STRUBBELIG. Development **148**

**Czechowski T, Stitt M, Altmann T, Udvardi MK, Scheible W-Rd** (2005) Genome-Wide Identification and Testing of Superior Reference Genes for Transcript Normalization in Arabidopsis. Plant Physiology **139:** 5-17

**Engelsdorf T, Gigli-Bisceglia N, Veerabagu M, McKenna JF, Vaahtera L, Augstein F, Van der Does D, Zipfel C, Hamann T** (2018) The plant cell wall integrity maintenance and immune signaling systems cooperate to control stress responses in *Arabidopsis thaliana*. Science Signaling **11**

**Giovannoni M, Lironi D, Marti L, Paparella C, Vecchi V, Gust AA, De Lorenzo G, Nürnberger T, Ferrari S** (2021) The Arabidopsis thaliana LysM-containing Receptor-Like Kinase 2 is required for elicitor-induced resistance to pathogens. Plant, Cell & Environment **44:** 3775-3792

**Glazebrook J, Ausubel FM** (1994) Isolation of phytoalexin-deficient mutants of *Arabidopsis thaliana* and characterization of their interactions with bacterial pathogens. Proceedings of the National Academy of Sciences **91:** 8955-8959

**Guo H, Li L, Ye H, Yu X, Algreen A, Yin Y** (2009) Three related receptor-like kinases are required for optimal cell elongation in *Arabidopsis thaliana*. Proceedings of the National Academy of Sciences **106:** 7648-7653

**Hématy K, Sado P-E, Van Tuinen A, Rochange S, Desnos T, Balzergue S, Pelletier S, Renou J-P, Höfte H** (2007) A receptor-like kinase mediates the response of Arabidopsis cells to the inhibition of cellulose synthesis. Current Biology **17:** 922-931

**Liu Y, Zhang S** (2004) Phosphorylation of 1-Aminocyclopropane-1-Carboxylic Acid Synthase by MPK6, a Stress-Responsive Mitogen-Activated Protein Kinase, Induces Ethylene Biosynthesis in Arabidopsis. The Plant Cell **16:** 3386-3399

**Mao G, Meng X, Liu Y, Zheng Z, Chen Z, Zhang S** (2011) Phosphorylation of a WRKY Transcription Factor by Two Pathogen-Responsive MAPKs Drives Phytoalexin Biosynthesis in Arabidopsis. The Plant Cell **23:** 1639-1653

**Millet YA, Danna CH, Clay NK, Songnuan W, Simon MD, Werck-Reichhart D, Ausubel FM** (2010) Innate Immune Responses Activated in Arabidopsis Roots by Microbe-Associated Molecular Patterns The Plant Cell **22:** 973-990

**Müller TM, Böttcher C, Morbitzer R, Götz CC, Lehmann J, Lahaye T, Glawischnig E** (2015) TRANSCRIPTION ACTIVATOR-LIKE EFFECTOR NUCLEASE-Mediated Generation and Metabolic Analysis of Camalexin-Deficient *cyp71a12 cyp71a13* Double Knockout Lines. Plant Physiology **168:** 849-858

**Nafisi M, Goregaoker S, Botanga CJ, Glawischnig E, Olsen CE, Halkier BA, Glazebrook J** (2007) Arabidopsis Cytochrome P450 Monooxygenase 71A13 Catalyzes the Conversion of Indole-3-Acetaldoxime in Camalexin Synthesis. The Plant Cell **19:** 2039-2052

**Nakagawa Y, Katagiri T, Shinozaki K, Qi Z, Tatsumi H, Furuichi T, Kishigami A, Sokabe M, Kojima I, Sato S, Kato T, Tabata S, Iida K, Terashima A, Nakano M, Ikeda M, Yamanaka T, Iida H** (2007) Arabidopsis plasma membrane protein crucial for Ca2+ influx and touch sensing in roots. Proceedings of the National Academy of Sciences **104:** 3639-3644

**Park J-H, Halitschke R, Kim HB, Baldwin IT, Feldmann KA, Feyereisen R** (2002) A knock-out mutation in allene oxide synthase results in male sterility and defective wound signal transduction in Arabidopsis due to a block in jasmonic acid biosynthesis. The Plant Journal **31:** 1-12

**Renard J, Niñoles R, Martínez-Almonacid I, Gayubas B, Mateos-Fernández R, Bissoli G, Bueso E, Serrano R, Gadea J** (2020) Identification of novel seed longevity genes related to oxidative stress and seed coat by genome-wide association studies and reverse genetics. Plant, Cell & Environment **43:** 2523-2539

**Schmidt A, Mächtel R, Ammon A, Engelsdorf T, Schmitz J, Maurino VG, Voll LM** (2020) Reactive oxygen species dosage in Arabidopsis chloroplasts can improve resistance towards Colletotrichum higginsianum by the induction of WRKY33. New Phytologist **226:** 189-204

**Völkner C, Holzner LJ, Day PM, Ashok AD, Vries Jd, Bölter B, Kunz H-H** (2021) Two plastid POLLUX ion channel-like proteins are required for stress-triggered stromal Ca2+release. Plant Physiology **187:** 2110-2125

**Wang H, Ngwenyama N, Liu Y, Walker JC, Zhang S** (2007) Stomatal Development and Patterning Are Regulated by Environmentally Responsive Mitogen-Activated Protein Kinases in Arabidopsis. The Plant Cell **19:** 63-73

**Weinl S, Held K, Schlücking K, Steinhorst L, Kuhlgert S, Hippler M, Kudla J** (2008) A plastid protein crucial for Ca2+-regulated stomatal responses. New Phytologist **179:** 675-686

**Yamaguchi Y, Huffaker A, Bryan AC, Tax FE, Ryan CA** (2010) PEPR2 Is a Second Receptor for the Pep1 and Pep2 Peptides and Contributes to Defense Responses in Arabidopsis The Plant Cell **22:** 508-522

**Zhang J, Eswaran G, Alonso-Serra J, Kucukoglu M, Xiang J, Yang W, Elo A, Nieminen K, Damén T, Joung J-G, Yun J-Y, Lee J-H, Ragni L, Barbier de Reuille P, Ahnert SE, Lee J-Y, Mähönen AP, Helariutta Y** (2019) Transcriptional regulatory framework for vascular cambium development in Arabidopsis roots. Nature Plants **5:** 1033-1042
